# The RNA demethylase FTO targets m^6^Am in snRNA to establish distinct methyl isoforms that influence splicing

**DOI:** 10.1101/327924

**Authors:** Jan Mauer, Miriam Sindelar, Théo Guez, Jean-Jacques Vasseur, Andrea Rentmeister, Steven S. Gross, Livio Pellizzoni, Françoise Debart, Hani Goodarzi, Samie R. Jaffrey

## Abstract

Small nuclear RNAs (snRNAs) are core spliceosome components and mediate pre-mRNA splicing. During their biogenesis, snRNAs acquire several constitutive nucleotide modifications. Here we show that snRNAs also contain a regulated and reversible nucleotide modification causing them to exist as two different methyl isoforms, m_1_ and m_2_, reflecting the methylation state of the adenosine adjacent to the snRNA cap. We find that snRNA biogenesis involves the formation of an initial m_1_-isoform with a single-methylated adenosine (2’-*O*-methyladenosine, Am), which is then converted to a dimethylated m_2_-isoform (*N*^6^,2’-*O*-dimethyladenosine, m^6^Am). The relative m_1_- and m_2_-isoform levels are determined by the RNA demethylase FTO, which selectively demethylates the m_2_-isoform. We show FTO is inhibited by endogenous metabolites, resulting in increased m_2_-snRNA levels. Furthermore, cells that exhibit high m_2_-snRNA levels show altered patterns of alternative splicing. Together, these data reveal that FTO has a central role in snRNA biogenesis and controls a previously unknown step of snRNA processing involving reversible methylation, thereby providing a potential link between reversible RNA modifications and mRNA splicing.

---

Small nuclear RNAs (snRNAs) are among the most abundant and extensively studied RNAs in eukaryotic cells. These uridine (U)-rich non-coding RNAs – which include U1, U2, U4, U5, and U6 – were discovered nearly fifty years ago^1^, and studies of their function lead to the elucidation of the fundamental molecular mechanisms that mediate pre-mRNA splicing^2-5^.

snRNAs undergo a series of processing events that are required for incorporation into spliceosomes^6-8^. Except for U6 and U6atac snRNA^9^, snRNAs are synthesized by RNA polymerase II as precursors that initially contain a 3’-end extension and acquire an *N*-^7^- methylguanosine (m^7^G) cap^10^. These snRNAs are exported into the cytosol where they are incorporated into small nuclear ribonucleoproteins (snRNPs) by binding to Sm proteins^11^. Their m^7^G cap is then further methylated to form the *N*^2,2,7^-trimethylguanosine (TMG) cap and the snRNA 3’-end is trimmed^12-14^. snRNAs become highly stable as a result of their incorporation into snRNPs and are transported back to the nucleus where they assemble into spliceosomes that mediate pre-mRNA splicing^6-8^. snRNAs that are not properly processed are not incorporated into snRNPs and are therefore unstable and degraded^15,16^.

Importantly, snRNAs contain a set of constitutive nucleotide modifications that are highly conserved across species and essential for snRNA integrity and function^17,18^. Thus, all mature snRNAs are thought to exist as a single molecular species, and deviations outside of this species are not thought to be utilized by spliceosomes.

Here we show that all Sm-class spliceosomal snRNAs can exist as two distinct subtypes, differing by a single methyl modification. The transcription-start nucleotide of most snRNAs is an adenosine with a constitutive 2’-*O*-methyl modification on the ribose sugar. We find that these adenosines are subjected to a second, reversible methylation selectively at the *N*^6^-position on the adenine base. This methylation converts the canonical single-methylated m_1_-snRNA to a dimethylated m_2_-snRNA. m_2_-snRNAs are assembled into snRNPs and promote the splicing of exons that are normally poorly included in cells. Furthermore, we find that m_2_-snRNAs are major targets of the RNA demethylase FTO, which is highly selective for demethylating the *N*^6^- methyl group in m^6^Am. We find that FTO functions early during snRNA biogenesis to convert the m_2_-snRNA to m_1_-snRNA. FTO-mediated demethylation of m_2_-snRNA is blocked by intracellular metabolites, including the oncometabolite 2-hydroxyglutarate, linking metabolism to snRNA methylation state, FTO activity, and splicing. Taken together, these data reveal a methylation switch, controlled by FTO, that results in the formation of a distinct form of snRNAs that may influence transcriptome diversity.

## The RNA demethylase FTO selectively demethylates snRNAs

The concept that RNA can undergo reversible methylation was first proposed based on studies of fat mass and obesity-associated protein (FTO)^19^. FTO is a member of the ALKB family of DNA nucleotide repair enzymes, but was found to exhibit weak demethylase activity towards methylated RNA nucleotides such as *N*^3^-methyluridine (m^3^U) and *N*^6^-methyladenosine (m^6^A)^19,20^. We recently showed that FTO shows highly efficient demethylation of *N*^6^,2’-*O*dimethyladenosine (m^6^Am), with nearly 100-fold higher catalytic activity towards m^6^Am compared to m^6^A (ref. 21). FTO selectively demethylates the *N*^6^-methyl modification, resulting in 2’-*O*-methyladenosine (Am)^21^. m^6^Am is exclusively found within the “extended cap” of RNA polymerase II-transcribed RNAs, at the transcription-start nucleotide immediately adjacent to the m^7^G (ref. 22,23).

The physiological importance of RNA demethylation was demonstrated by analysis of FTO-deficient mice, which exhibit altered synaptic transmission, metabolic abnormalities, and growth retardation^24,25^. Additionally, humans with loss-of-function FTO mutations show severe developmental defects and microcephaly^26^. FTO has been proposed to mediate its effects by influencing alternative splicing of mRNA, based on transcriptome-wide splicing alterations detected in FTO-deficient cells^27,28^.

It remains unclear which methylated residues in the transcriptome are preferentially targeted by FTO. m^6^Am shows high stoichiometry in mRNA, which suggests that FTO does not efficiently demethylate m^6^Am to Am in mRNA^21^. Indeed, nearly all the m^6^Am peaks in mRNA showed increases of less than 10% in *Fto* knockout brain compared to wild-type^21^. Although FTO clearly demethylates m^6^Am in mRNA, the moderate effect of FTO-mediated mRNA demethylation likely reflects the fact that FTO is nuclear, while mRNAs are predominantly cytosolic^21^. Although FTO can translocate to the cytoplasm and have access to mRNA under certain stress conditions, such as nutrient deprivation^29^, in most cells and tissues, FTO is detected exclusively in the nucleus^19,25^. Thus, FTO may have nuclear RNA targets that might account for the diverse phenotypes observed in FTO-deficient mice and humans.

To identify the RNAs that are under control of FTO, we performed miCLIP analysis to map *N*^6^- methyladenine (6mA) in wild-type and *Fto*-knockout mouse liver RNA (**Extended Data Fig. 1a**). Unlike in previous studies^24,25^, no poly(A) purification was used, thus allowing all cellular RNAs to be analyzed. To detect m^6^Am, we searched for RNA species that show increased methylation at the transcription-start nucleotide in the *Fto* knockout transcriptome compared to wild type. This analysis revealed only a small subset of methylation events that were significantly increased in the *Fto* knockout transcriptome (**Fig. 1a**). Examination of these RNAs showed that they were exclusively small nuclear RNAs (snRNAs), including U1, U4atac, and U5 (**Fig. 1a**).

**Figure 1.**
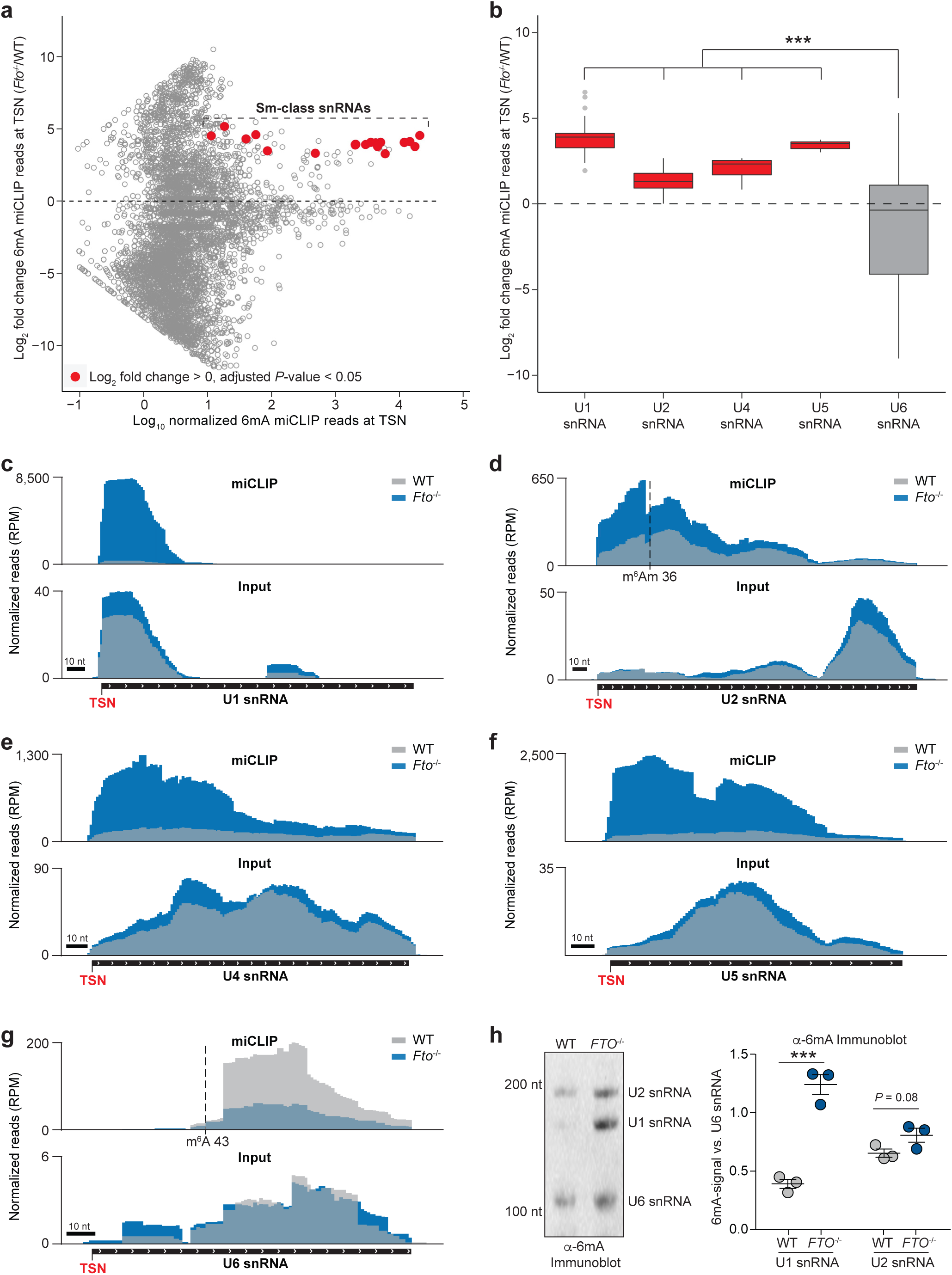
FTO selectively demethylates small nuclear RNAs. **a**, *N*^6^-methyladenine (6mA) mapping in total RNA using miCLIP reveals FTO-dependent demethylation of small nuclear RNAs (snRNAs). To measure FTO-regulated m^6^Am sites in RNAs, 6mA miCLIP reads were counted in a 20-nucleotide window surrounding the transcription-start nucleotide (TSN) and normalized to input RNA-Seq reads in the same region. The relative expression changes were calculated. The MA plot shows the log_2_ fold change in methylation in *Fto* knockout mouse liver (*Fto*^-/-^) compared to wild-type liver (WT). Transcripts that showed statistically significant upregulation of transcription-start nucleotide methylation are indicated in red. These transcripts are derived exclusively from small nuclear RNA genes. Data represents the average from datasets of three independent biological replicates per genotype. **b,** FTO deficiency leads to increased transcription-start nucleotide methylation of major spliceosomal snRNAs. miCLIP reads were counted in a 20-nucleotide window surrounding the transcription-start nucleotide, normalized to input RNA-Seq reads in the same region and relative expression changes were calculated. In this analysis, no *P*-value cut-off was applied. Instead, the mean log_2_-fold change in transcription-start nucleotide methylation of specific snRNA gene classes (U1, U2, U4, U5 and U6) in *Fto* knockout mouse liver (*Fto*^-/-^) compared to wild-type liver (WT) is shown. Notably, only snRNA genes transcribed by RNA polymerase II increased methylation, whereas U6 snRNA genes (grey), which are transcribed by RNA polymerase III and thus do not acquire a m^7^G cap, show no change in transcription-start nucleotide methylation upon FTO deficiency (Data represents the average from datasets of three independent biological replicates per genotype; one-way ANOVA with Tukey’s post hoc test \*\*\**P* ≤ 0.001). **c**, U1 snRNA shows increased transcription-start nucleotide methylation in FTO-deficient mouse liver. The grey tracks denote WT liver, whereas the blue track denotes *Fto*^*-/-*^liver. A representative read coverage track for U1 snRNA is shown. 6mA miCLIP reads are shown in the upper panel. As can be seen, miCLIP reads are readily detected at the transcription-start nucleotide (TSN) in the U1 snRNA in *Fto*^-/-^RNA, but is nearly undetectable in U1 snRNA from wild-type RNA. Input reads are shown in the lower panel. RPM = reads per million mapped reads; data represents the combined tracks from datasets of three independent biological replicates per genotype. **d**, U2 snRNA shows increased transcription-start nucleotide methylation in FTO-deficient mouse liver. The grey tracks denote WT liver, whereas the blue track denotes *Fto*^*-/-*^liver. A representative read coverage track for U2 snRNA is shown. 6mA miCLIP reads are shown in the upper panel. As can be seen, miCLIP reads are substantially higher at the transcription-start nucleotide (TSN) in the U2 snRNA in *Fto*^-/-^RNA, but is reduced in the U2 snRNA from wildtype RNA. Input reads are shown in the lower panel. The transcription-start nucleotide is indicated in red and the constitutive internal *N*^6^,2’-*O*-dimethyladenosine (m^6^Am) residue at position 36 is indicated by the black dashed line. (RPM = reads per million mapped reads; data represents the combined tracks from datasets of three independent biological replicates per genotype). **e**, U4 snRNA shows increased transcription-start nucleotide methylation in FTO-deficient mouse liver. The grey tracks denote WT liver, whereas the blue track denotes *Fto*^*-/-*^liver. A representative read coverage track for U4 snRNA is shown. 6mA miCLIP reads are shown in the upper panel. As can be seen, miCLIP reads are substantially higher at the transcription-start nucleotide (TSN) in the U4 snRNA in *Fto*^-/-^RNA, but is reduced in the U4 snRNA from wildtype RNA. Input reads are shown in the lower panel. The transcription-start nucleotide is indicated in red (RPM = reads per million mapped reads; data represents the combined tracks from datasets of three independent biological replicates per genotype). **f**, U5 snRNA shows increased transcription-start nucleotide methylation in FTO-deficient mouse liver. The grey tracks denote WT liver, whereas the blue track denotes *Fto*^-/-^liver. A representative read coverage track for U5 snRNA is shown. 6mA miCLIP reads are shown in the upper panel. As can be seen, miCLIP reads are substantially higher at the transcription-start nucleotide (TSN) in the U5 snRNA in *Fto*^*-/-*^RNA, but is reduced in the U5 snRNA from wildtype RNA. Input reads are shown in the lower panel. The transcription-start nucleotide is indicated in red (RPM = reads per million mapped reads; data represents the combined tracks from datasets of three independent biological replicates per genotype). **g**, U6 snRNA transcription-start nucleotide methylation is not affected in FTO-deficient mouse liver. The grey tracks denote WT liver, whereas the blue track denotes *Fto*^*-/-*^liver. A representative read coverage track for U6 snRNA is shown. 6mA miCLIP reads are shown in the upper panel. As can be seen, there are essentially no detectable miCLIP reads at the TSN in either the *Fto*^*-/-*^or wildtype miCLIP samples in the U6 RNA. Therefore, U6 snRNA lacks 6mA at the TSN. Input reads are shown in the lower panel. The constitutive internal *N*^6^methyladenosine (m^6^A) residue at position 43 is indicated by the black dashed line. RPM = reads per million mapped reads; data represents the combined tracks from datasets of three independent biological replicates per genotype. **h**, FTO deficiency leads to increased methylation of mature snRNAs. 6mA-immunoblot of wildtype (WT) and FTO knockout (*FTO*^-/-^) HEK293 cells. Methylation of the most abundant snRNAs (U1, U2, U6) was detected with an antibody directed against 6mA. The molecular weights of the 6mA-reactive bands are consistent with the mature processed forms of the snRNA, demonstrating that the mature snRNA, rather than an snRNA precursor, contains elevated 6mA in *FTO* knockout cells. The left panel shows a representative 6mA-immunoblot with the positions of nucleotide size markers and the individual snRNAs indicated on the left and right, respectively. The right panel shows the quantification of 6mA signal in U1 and U2 snRNA relative to the 6mA signal in U6 snRNA. U6 snRNA was used for normalization since it has an internal m^6^A residue that is constitutive and lacks m^6^Am, and therefore is not regulated by FTO. Notably, we observed strong 6mA immunoreactivity towards U2 snRNA in WT cells and increased methylation in *Fto*^*-/-*^cells did not reach statistical significance. However, U2 snRNA contains a constitutive internal m^6^Am residue (see also **Fig. 1d**) that is not demethylated by FTO, which likely masks FTO deficiency-dependent effects (*n* = 3 independent biological replicates; mean ± s.e.m.; unpaired student’s *t*-test \*\*\**P* ≤ 0.001).

To test whether increased transcription-start nucleotide methylation in the *Fto* knockout transcriptome is observed in all spliceosomal snRNAs, we next looked specifically at the methylation fold change of Sm- and Lsm-class snRNAs. We observed increased transcription-start nucleotide methylation of all Sm-class snRNAs of the major and minor spliceosome (**Fig. 1b, Extended Data Fig. 1b**). In each case, the 6mA reads were detected around the transcription-start nucleotide and were between 10-20-fold higher in the *Fto* knockout compared to the wild-type transcriptome (**Figs. 1c-f, Extended Data Figs. 2a-c**). Moreover, U7 snRNA, which is involved in 3’-end processing of histone mRNAs^30^ and the small nucleolar RNAs (snoRNAs) U3 and U8 that function in rRNA processing also showed increases transcription-start nucleotide methylation in *Fto* knockout compared to wild-type transcriptome (**Extended Data Figs. 3a-c**).

The increase in 6mA reads was seen in all Sm-class snRNAs, but was not seen in the Lsm-class snRNAs U6 and U6atac – the only two spliceosomal snRNAs that are transcribed by RNA polymerase III rather than RNA polymerase II (ref. 9) (**Figs. 1b, g and Extended Data Figs. 1b, 2d**). U6 and U6atac do not acquire an m^7^G cap; however, U6 snRNA contains an internal m^6^A at position 43 (ref. 17). Nevertheless, FTO depletion did not lead to significant upregulation of 6mA reads at this internal position (**Figs. 1b, g**). Similarly, U2 snRNA, which contains an m^6^Am at position 36 (ref. 17), did not show a clear increase of 6mA reads at this internal position (**Figs. 1d, g**). Thus, the FTO-regulated sites are specifically localized to the transcription-start nucleotide region of Sm-class snRNAs.

miCLIP does not readily distinguish between mature snRNA and the unprocessed longer snRNA precursors or truncated forms. Therefore, we asked if the increased snRNA methylation upon *FTO* knockout could be detected in mature snRNAs. To do this, we used 6mA immunoblotting to detect the length of 6mA-immunoreactive snRNAs. In these experiments, we performed 6mA immunoblot of small RNA (<200 nt) from wild-type and *FTO* knockout HEK293 cells. We observed bands corresponding to the mature snRNA forms, with significantly increased 6mAimmunoreactivity in *FTO* knockout cells (**Fig. 1h**).

Taken together, these data indicate that all Sm-class snRNAs are substrates for FTO-mediated demethylation during their life cycle.

## The first nucleotide of snRNAs is reversibly methylated

The finding that Sm-class snRNA modification is regulated by FTO is surprising, since these snRNAs are not known to contain m^6^Am or m^6^A at the transcription-start nucleotide. Instead, Sm-class snRNAs are thought to only contain 2’-*O*-methyladenosine (Am) at this position^13^. To biochemically determine the identity of the first nucleotide in snRNAs in wild-type and *FTO* knockout cells, we used a thin-layer chromatography (TLC)-based approach^23^. In this assay, the 5’-cap is removed from RNA, followed by radiolabeling of the exposed 5’ nucleotide with [*γ*^32^P]-ATP. Next, the snRNAs are treated with ribonuclease and the nucleotide hydrolysate is resolved by 2D-TLC. This approach readily resolves whether the first nucleotide is m^6^Am or Am, or any other nucleotide based on its migration pattern^23^.

We first purified the small RNA fraction (<200 nt), since this fraction is highly enriched in snRNAs. Quantification of the first nucleotide of these RNAs from wild-type cells showed a high level of Am, as well as low, but measurable levels of m^6^Am (**Fig. 2a**). However, when the small RNA fraction was prepared from *FTO* knockout HEK293 cells, a substantial increase in the level of m^6^Am at the first nucleotide was readily detected (**Fig. 2a**).

**Figure 2.**
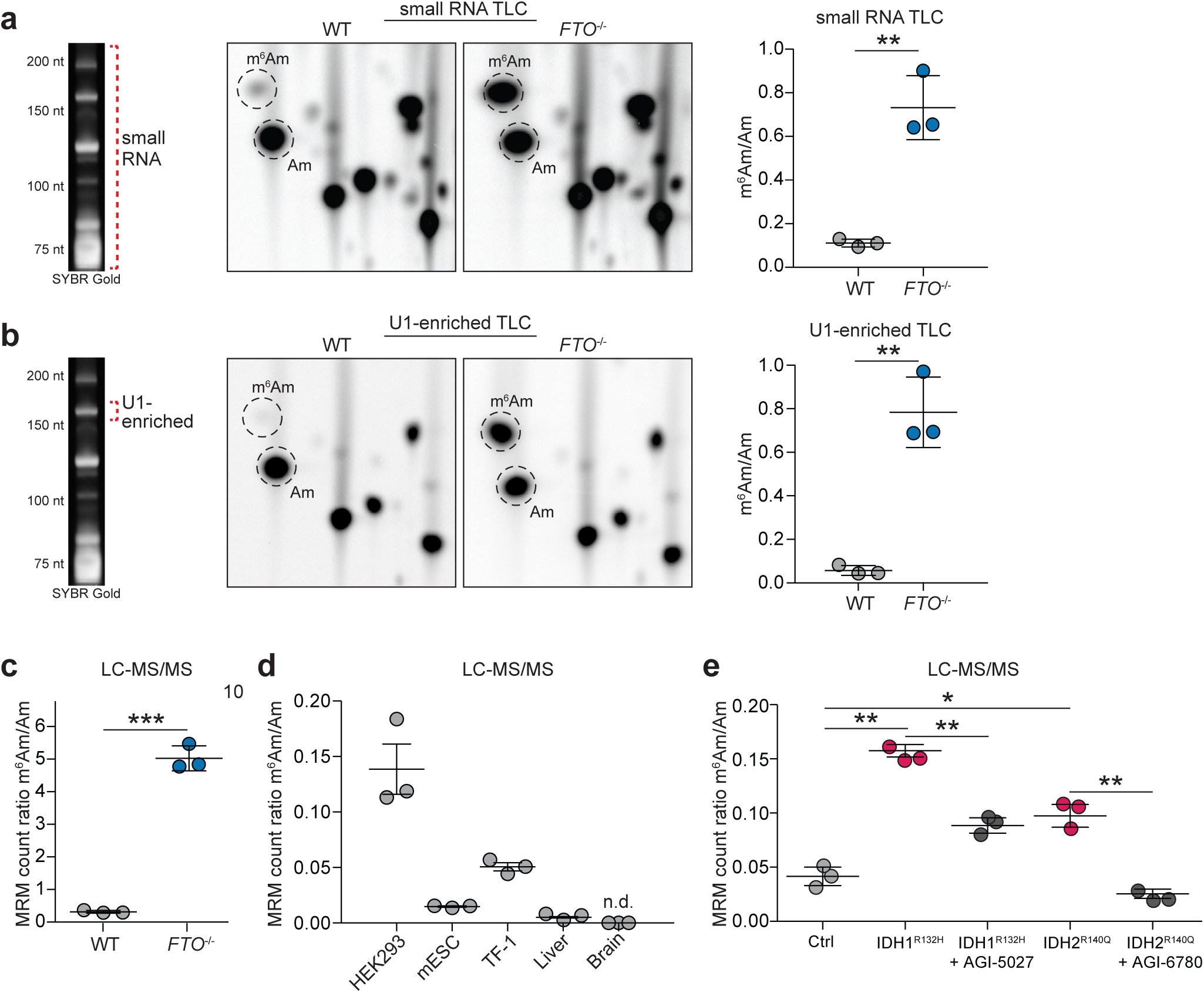
Reversible *N*^6^,2’-*O*-dimethyladenosine (m^6^Am) in small nuclear RNAs. **a**, FTO deficiency reveals the presence of m^6^Am in small RNAs. The relative abundance of modified adenosines in small RNA caps derived from wild-type (WT) and FTO-deficient (*FTO*^-/-^) HEK293 cells was determined by thin layer chromatography. The left panel shows a TBE-Urea gel image stained with SYBR Gold, where the red dashed line indicates the small RNA fraction (all RNAs < 200 nt) that was used for the analysis. The middle panel shows a representative image of the migration pattern of radiolabeled nucleotides, where the position of m^6^Am and 2’*O*-methyladenosine (Am) is indicated by the dashed black circles. The level of m^6^Am is markedly increased in small RNA in *FTO* knockout cells. The right panel shows the quantification of the m^6^Am/Am ratio in small RNA (*n* = 3 independent biological replicates; mean ± s.d; unpaired student’s *t*-test \*\**P* ≤ 0.01). **b**, FTO deficiency reveals the presence of m^6^Am in U1 snRNA. In contrast to (**a**), here we isolated the U1 snRNA by excising the RNA from a TBE-Urea gel image stained with SYBR Gold (left). The red dashed line indicates the U1-enriched fraction that was gel-extracted and used for the analysis. The relative abundance of m^6^Am and Am in U1 snRNA caps derived from wild-type (WT) and FTO-deficient (*FTO*^-/-^) HEK293 cells was determined by thin layer chromatography. The level of the m^6^Am cap in U1 snRNA is markedly increased in *FTO* knockout cells. The migration position of m^6^Am and 2’-*O*-methyladenosine (Am) is indicated by the dashed black circles. The right panel shows the quantification of the m^6^Am/Am ratio in the U1-enriched fraction (*n* = 3 independent biological replicates; mean ± s.d; unpaired student’s *t*-test \*\**P* ≤ 0.01). **c**, FTO deficiency increases the relative abundance of m_2_-snRNA caps in HEK293 cells. Small RNA from WT and *Fto*^*-/-*^was digested with nuclease P1 to specifically liberate the extended cap structure dinucleotide and analyzed by LC-MS/MS. Shown is the ratio of m_2_-snRNA caps (m^6^Am) to m_1_-snRNA caps (Am) represented by the integrated peak area ratio of the corresponding MRM transitions (recorded in positive ion mode) (n=3 independent biological replicates, mean ± s.d.; unpaired Student’s t-test, *** p < 0.001). **d**, Differential expression of m_2_-snRNA (cap-ppp-m^6^Am) caps across cells and tissues. Small RNA from wild-type HEK293 cells, naïve mouse embryonic stem cells (mESCs), TF-1 erythroleukemia cells, as well as mouse liver and brain tissue was digested with nuclease P1 to specifically liberate the extended cap structure dinucleotide and analyzed by LC-MS/MS. Shown is the ratio of m_2_-snRNA caps (m^6^Am) to m_1_-snRNA caps (Am) represented by the integrated peak area ratio of the corresponding MRM transitions (recorded in positive ion mode). These data suggest that the abundance of m_2_-snRNAs varies in a cell-specific manner (n.d.=cap-pppm^6^Am not detected; n=3 independent biological replicates, mean ± s.d.; unpaired Student’s t-test,** p < 0.01). **e**, Increased abundance of m_2_-snRNA caps in oncometabolite-dependent cancer. Small RNA from wild-type (Ctrl) and mutant IDH-expressing cells was digested with nuclease P1 to specifically liberate the extended cap structure dinucleotide and analyzed by LC-MS/MS. Shown is the ratio of m_2_-snRNA caps (m^6^Am) to m_1_-snRNA caps (Am) represented by the integrated peak area ratio of the MRM transition (recorded in positive ion mode). IDH1^R132H^and IDH2^R140Q^-expressing TF-1 cells have high levels of 2-hydroxyglutarate (2-HG). 2-HG is a natural inhibitor of FTO activity^32^ and leads to increased abundance of m_2_-snRNAs. Specific inhibition of the mutant IDH1 (AGI-5027) and IDH2 (AGI-6780) isoforms shows that these effects are reversible by blocking production of 2HG (n=3 independent biological replicates, mean ± s.d.; one-way ANOVA with Tukey’s post hoc test \**P* ≤ 0.05, \*\**P* ≤ 0.01).

To ensure that FTO depletion is affecting snRNA and not some other type of small RNA, we directly measured the first nucleotide in U1 snRNA. To do this, we gel-purified U1 snRNA, a 164 nucleotide-long snRNA that migrates to a well-defined position by PAGE (**Figs. 1f, 2b**). We then used the TLC approach to determine the first nucleotide in the U1 snRNA-enriched fraction. Consistent with previous publications^13^, the first nucleotide in U1 was predominantly Am in wild-type HEK293 cells (**Fig. 2b**). However, in U1 snRNA from *FTO* knockout HEK293 cells, m^6^Am was markedly increased, with levels similar to Am (**Fig. 2b**). These data indicate that snRNAs can contain m^6^Am at the first encoded nucleotide.

Among the snRNAs, U2 showed the smallest increase in the miCLIP signal in *FTO* knockout cells. U2 is difficult to analyze since it contains a constitutive internal m^6^Am, which causes this snRNA to be immunoprecipitated irrespective of the methylation status of its transcription-start nucleotide (**Fig. 1d**). Therefore, we also analyzed the 5’-nucleotide of this snRNA by TLC. Indeed, we observed that *FTO* knockout resulted in a substantial increase in m^6^Am at the 5’position of U2 snRNA (**Extended Data Fig. 4a**). This indicates that U2 also contains an m^6^Am nucleotide that is regulated by FTO.

To provide an independent measurement to assess whether snRNAs can contain m^6^Am at the first nucleotide, we developed a mass spectrometry-based assay. To ensure that we examined only m^6^Am residues in the context of the cap structure, small RNA was treated with P1 nuclease, resulting in all internal nucleotides being digested to mononucleotides. However, under these conditions, the first nucleotide remains connected to the cap in the form of a “cap-dinucleotide” structure (cap-ppp-Am or cap-ppp-m^6^Am) (**Extended Data Fig. 5a**).

Cap-dinucleotides were readily quantified by high-resolution mass spectrometry analysis using positive ion mode detection, since negative ion mode did not allow sensitive detection of cap-dinucleotides (**Extended Data Fig. 5b**). A multiple reaction monitoring protocol was developed based on the fragment ion transitions from distinct dinucleotide precursor species (**Extended Data Fig. 5c**). This approach exhibited high sensitivity and linear detection of the first nucleotide within the cap context (**Extended Data Fig. 5d**).

Using this approach, we verified that Am was associated with the cap after digestion of small RNA from wild-type HEK293 cells (**Fig. 2c, d**). However, m^6^Am was the predominant form in *FTO* knockout HEK293 cells, with a 6.3-fold enrichment of m^6^Am over Am (**Fig. 2c**). Thus, both the TLC and mass spectrometry analysis indicate that snRNAs contain m^6^Am at the first nucleotide in FTO-deficient cells.

Together, these results suggest that during snRNA biogenesis, snRNAs transiently exist as a dimethylated form (m_2_-snRNA) with m^6^Am as the first nucleotide. The m_2_-snRNA can then be demethylated by FTO to a single-methylated snRNA (m_1_-snRNA) with Am as the first nucleotide. The m_2_-snRNAs are readily detectable in HEK293 cells when FTO is depleted.

## m_2_-snRNAs are detected in different cells and tissues

The FTO depletion studies suggest that m_2_-snRNAs are efficiently demethylated during snRNA biogenesis in HEK293 cells. However, it is not clear if other cell types normally accumulate m_2_ snRNAs. As a first test, we measured m_1_-snRNAs and m_2_-snRNAs in different cell and tissue types. In these experiments, we isolated small RNAs and assayed the transcription-start nucleotide by mass spectrometry. In brain, only m_1_-snRNAs were detected (**Fig. 2d**). However, in liver, HEK293 cells, mouse embryonic stem cells (mESCs) and TF-1 erythroleukemia cells, m_2_-snRNAs were readily detected at 1-15% of the level of m_1_-snRNAs (**Fig. 2d** and **Extended Data Figs. 6a-d**). These data indicate that both m_1_-snRNAs and m_2_-snRNAs can exist in various cell and tissue types.

We next asked if m_1_-snRNAs and m_2_-snRNAs are regulated during embryonic stem cell development. To test this, we cultured mESCs for five days in the presence or absence of differentiation inhibitors. Mass spectrometry revealed that depletion of differentiation inhibitors induced a two-fold increase in m_2_-snRNAs compared to naive cells (**Extended Data Fig. 6b, c**).

Together, these data show that the levels of m_2_-snRNAs are regulated and expressed in a cell type-specific manner.

## m_2_-snRNAs are increased in response to the oncometabolite 2-hydroxyglutarate

Notably, FTO is inhibited by natural metabolites and oncometabolites that accumulate in cells under various developmental and disease states, including citrate, succinate, fumarate, and 2hydroxyglutarate (2-HG)^31,32^. These metabolites compete with *α*-ketoglutarate, an FTO cosubstrate for demethylation^31^. In the case of 2-HG, elevated levels are seen in cells expressing a cancer-associated mutant isocitrate dehydrogenase (IDH) variant^33^. Thus, we next asked if m_2_-snRNA levels are influenced by cellular metabolic states.

To determine if 2-HG affects snRNA methylation, we first measured m_1_-snRNAs and m_2_-snRNAs levels in small RNA isolated from TF-1 cells expressing the cancer-associated IDH1^R132H^ and IDH2^R140Q^ mutants^34,35^. Expression of mutant IDH2^R140Q^ elicited a marked increase in 2-HG levels when expressed in TF-1 cells, with a stronger effect seen with mutant IDH1^R132H^ (**Extended Data Fig. 7a**). In control cells, m_1_-snRNAs predominated as the major first nucleotide in small RNA. However, m_2_-snRNAs were markedly elevated following expression of both IDH1^R132H^ and IDH2^R140Q^, with a greater effect seen with IDH1^R132H^ (**Fig. 2e** and **Extended Data Fig. 7b**). Similar effects were seen using TLC analysis of U1 snRNA (**Extended Data Fig. 7c**).

To confirm that the effect of mutant IDH1 and IDH2 overexpression reflects increased 2-HG levels, we used specific inhibitors for each mutant isoform. Treatment with these inhibitors resulted in marked reduction of m_2_-snRNAs (**Fig. 2e** and **Extended Data Figs. 7b, c**).

Taken together, these data indicate that m_2_-snRNA levels can be altered by cellular metabolites such as 2-HG. Our finding suggests that m_2_-snRNAs have a role in 2-HG-associated pathogenic conditions.

## m^6^Am is demethylated prior to cap hypermethylation during snRNA biogenesis

The biogenesis of snRNAs has two nuclear phases^6-8^. The first nuclear phase comprises transcription and m^7^G capping of precursor snRNAs, which are then exported into the cytoplasm^8^. In the cytoplasm, the snRNAs acquire a TMG cap, and are then subjected to 3’ end trimming and nuclear re-import, all of which are dependent on the assembly of Sm proteins^6-8,14^. After the snRNAs are incorporated into small nuclear ribonucleoproteins (snRNPs), the second nuclear phase involves their function in splicing. Since FTO is found exclusively in the nucleus, the conversion of m_2_-snRNAs to m_1_-snRNAs by FTO likely takes place during one of these two nuclear phases. The absence or presence of a TMG cap is indicative of the specific nuclear phase of snRNAs.

Thus, we next asked if FTO demethylates m_2_-snRNAs before or after the formation of the TMG cap. We previously measured the demethylase activity of FTO towards m^6^Am using RNAs with an m^7^G cap^21^. We found the m^7^G cap and the *N*^7^-methyl modification were required for efficient demethylation^21^. To determine in which phase of snRNA biogenesis FTO acts, we chemically synthesized RNA substrates mimicking the pre-export and post-import cap structure of snRNA. These 20-nucleotide long RNAs started with either an m^7^G or a TMG cap, followed by a triphosphate linker, and m^6^Am. As we observed previously^21^, FTO treatment of m^7^G-capped RNA resulted in efficient demethylation of m^6^Am, as measured by the formation of Am (**Fig. 3a**). However, no Am formation was detected after FTO treatment of the post-import substrate comprising m^6^Am RNA and a TMG cap (**Fig. 3a**). These data suggest that m_2_-snRNAs are not a substrate for FTO after TMG cap formation.

**Figure 3.**
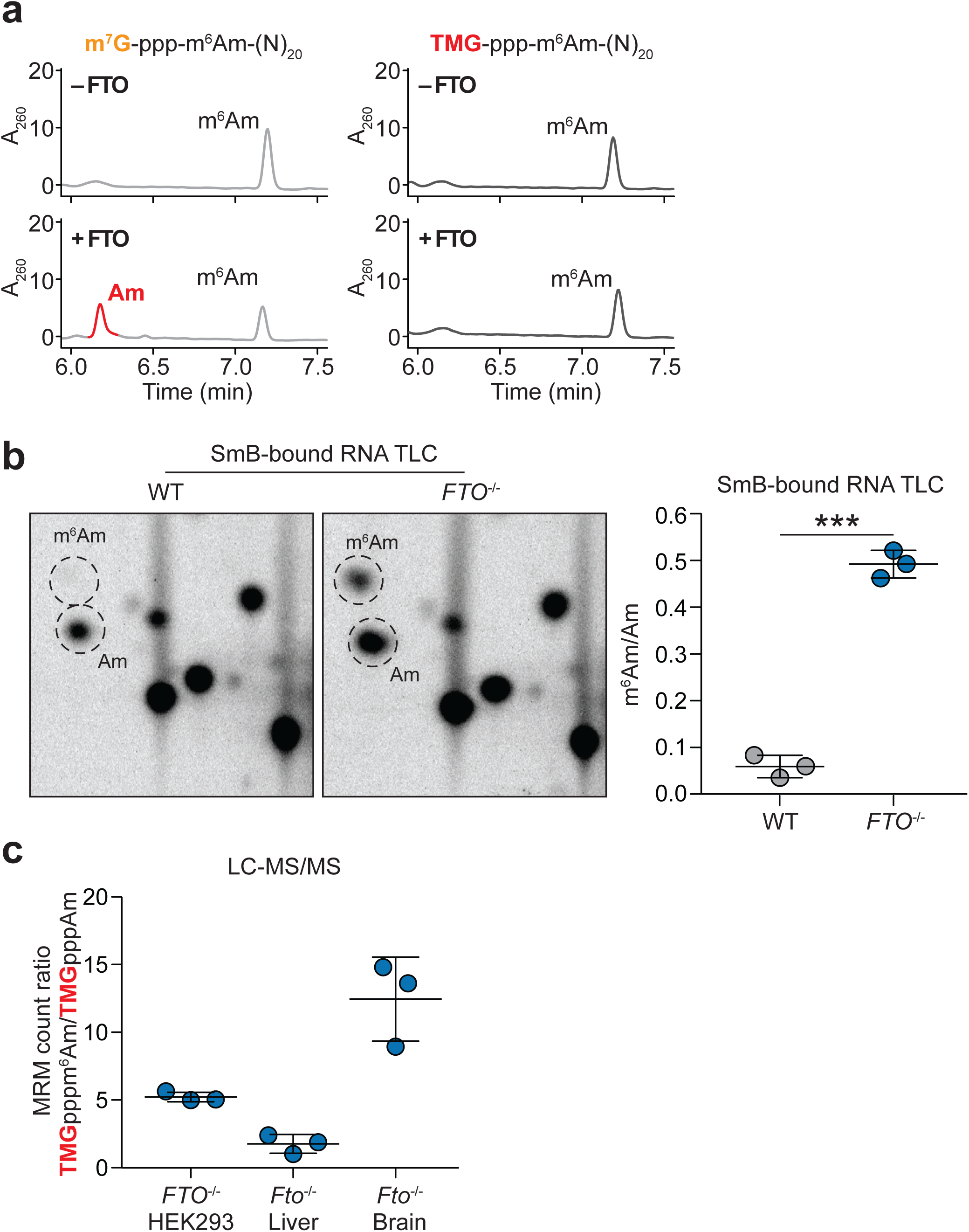
m2-snRNAs are incorporated into snRNPs. **a,** FTO does not exhibit measurable demethylation of m^6^Am in RNA containing a *N*^2,2,7^trimethylguanosine (TMG) cap. snRNAs contain a m^7^G cap shortly after their synthesis, but acquire a TMG cap during snRNA maturation. To determine whether FTO demethylates mature snRNA, we incubated full-length human FTO with synthetic 20-mer RNA oligonucleotides starting either with a 5’-m^7^G-ppp-m^6^Am or 5’-TMG-ppp-m^6^Am. In the context of an m^7^G cap, FTO readily converted m^6^Am to Am (left panels). However, the presence of an TMG cap completely blocked FTO demethylation activity towards m^6^Am (right panels), indicating that snRNAs starting with a TMG-ppp-m^6^Am are not a physiological target of FTO. This finding suggests that FTO demethylates m^6^Am in snRNA before the formation of an TMG cap (representative HPLC track shown; *n* = 3 independent experiments). **b**, m_2_-snRNAs are incorporated into small nuclear ribonucleoproteins (snRNPs). snRNA function is linked to its incorporation into spliceosomal snRNPs. To test whether m_2_-snRNAs are incorporated into snRNPs, immunoprecipitation of the SmB spliceosomal protein was performed. The relative abundance of modified adenosines in SmB-bound small RNA caps derived from wild-type (WT) and FTO-deficient (*FTO*^-/-^) HEK293 cells was determined by thin layer chromatography. In WT cells, SmB-bound carried predominately Am caps. However, in *Fto*^*-/-*^cells, m^6^Am was readily detected in the SmB-bound fraction, indicating that m_2_-snRNAs are incorporated into snRNPs and thus become part of the cellular splicing machinery. The left panel shows a representative image of the migration pattern of radiolabeled nucleotides, where the position of m^6^Am and Am is indicated by the dashed black circles. The right panel shows the quantification of the m^6^Am/Am ratio in small RNA (*n* = 3 independent biological replicates; mean ± s.d; unpaired student’s *t*-test \*\*\**P* ≤ 0.001). **c**, m_2_-snRNAs have TMG caps. We asked if the presence of m^6^Am in snRNA blocks the maturation of the snRNA cap from m^7^G to TMG. Small RNA from FTO-deficient HEK293 cells and tissues was digested with nuclease P1 to specifically liberate the extended cap structure dinucleotide and analyzed by LC-MS/MS. Small RNA from FTO-deficient cells exhibit high levels of m^6^Am, thereby allowing an assessment of whether m6Am affects the formation of TMG caps. Shown is the ratio of m_2_-snRNA TMG caps to m_1_-snRNA TMG caps represented by the integrated peak area ratio of the multiple reaction monitoring (MRM) transition 843.2 →→194.1 for TMG-ppp-m^6^Am and 829.1 → 194.1 for TMG-ppp-Am (recorded in positive ion mode). These data demonstrate that m_2_-snRNA contain TMG caps, thus indicating that m^6^Am does not function to regulate cap maturation (n=3 independent biological replicates, mean ± s.d.).

Based on the m^7^G cap-specific demethylation and considering the nuclear localization of FTO, we infer the decision to demethylate m_2_-snRNAs and form m_1_-snRNAs is determined during a specific, early phase of snRNA biogenesis. This phase occurs in the nucleus and the cytoplasm shortly after snRNA transcription but prior to the cytoplasmic formation of the TMG cap and assembly of snRNAs into snRNPs.

## m_2_-snRNAs acquire a TMG cap and assemble into snRNPs

In order to understand if m_2_-snRNAs could potentially function in splicing, we wanted to determine if m_2_-snRNAs assemble with Sm proteins into snRNPs. To test this, we immunoprecipitated snRNPs from cellular extracts using a small ribonucleoprotein particle protein B (SmB)-specific antibody. This approach has previously been shown to immunoprecipitate assembled snRNPs^36,37^. We then determined the first nucleotide of snRNAs in cellular snRNPs to determine the levels of m_1_-snRNAs and m_2_-snRNAs. As expected, in control HEK293 cells, predominantly m_1_-snRNAs were detected (**Fig. 3b**). However, in *FTO* knockout HEK293 cells, m_2_-snRNAs were substantially increased (**Fig. 3b**). These data indicate that m_2_-snRNAs assemble and form a distinct m^6^Am-containing class of snRNPs, i.e. m_2_-snRNPs.

After incorporation of snRNAs into snRNPs, their m^7^G cap is modified to form the TMG cap^14^. We asked if m_2_-snRNAs are similarly methylated to form the TMG cap. To test this, we measured the type of cap in m_2_-snRNAs by mass spectrometry. The small RNA fraction was purified from *FTO* knockout HEK293 cells and *Fto*-knockout mouse liver and brain. The RNA was digested with nuclease P1 to liberate the cap and m^6^Am as the dinucleotide cap structure (cap-ppp-m^6^Am). The presence of m^7^G or TMG was then assessed in the liberated cap structure. In each case, the primary cap structure was the TMG cap attached to m^6^Am (TMG-ppp-m^6^Am) (**Fig. 3c**, **Extended Data Fig. 6a**). This indicates that m_2_-snRNAs are subjected to cap hypermethylation during their biogenesis.

Taken together, these data show that m_2_-snRNAs undergo normal biogenesis, including incorporation into snRNPs and acquisition of the TMG cap structure.

## Altered splicing in cells expressing high levels of m_2_-snRNAs

As previously reported^28^, we found that the most pronounced effect of FTO deficiency is increased exon inclusion (**Fig. 4a**). To further refine this analysis, we next characterized the included exons for binding sites of known splicing factors in a previously published RNA-Seq analysis using control and FTO knockout HEK293 cells^27^. Here, we observed a significant and specific enrichment for SRSF1, HNRNPH2 and HNRNPK-dependent exons, whereas enrichment of other exons was not detected (**Fig. 4b** and **Extended Data Fig. 8a**). Similar effects were also seen in our independent HEK293T *FTO* knockdown RNA-Seq dataset (**Extended Data Figure 8b**). Together, these data indicate that an increase in m_2_-snRNPs is correlated with specific alternative splicing events, most notably inclusion of exons that would normally be inefficiently spliced into mature mRNA.

**Figure 4.**
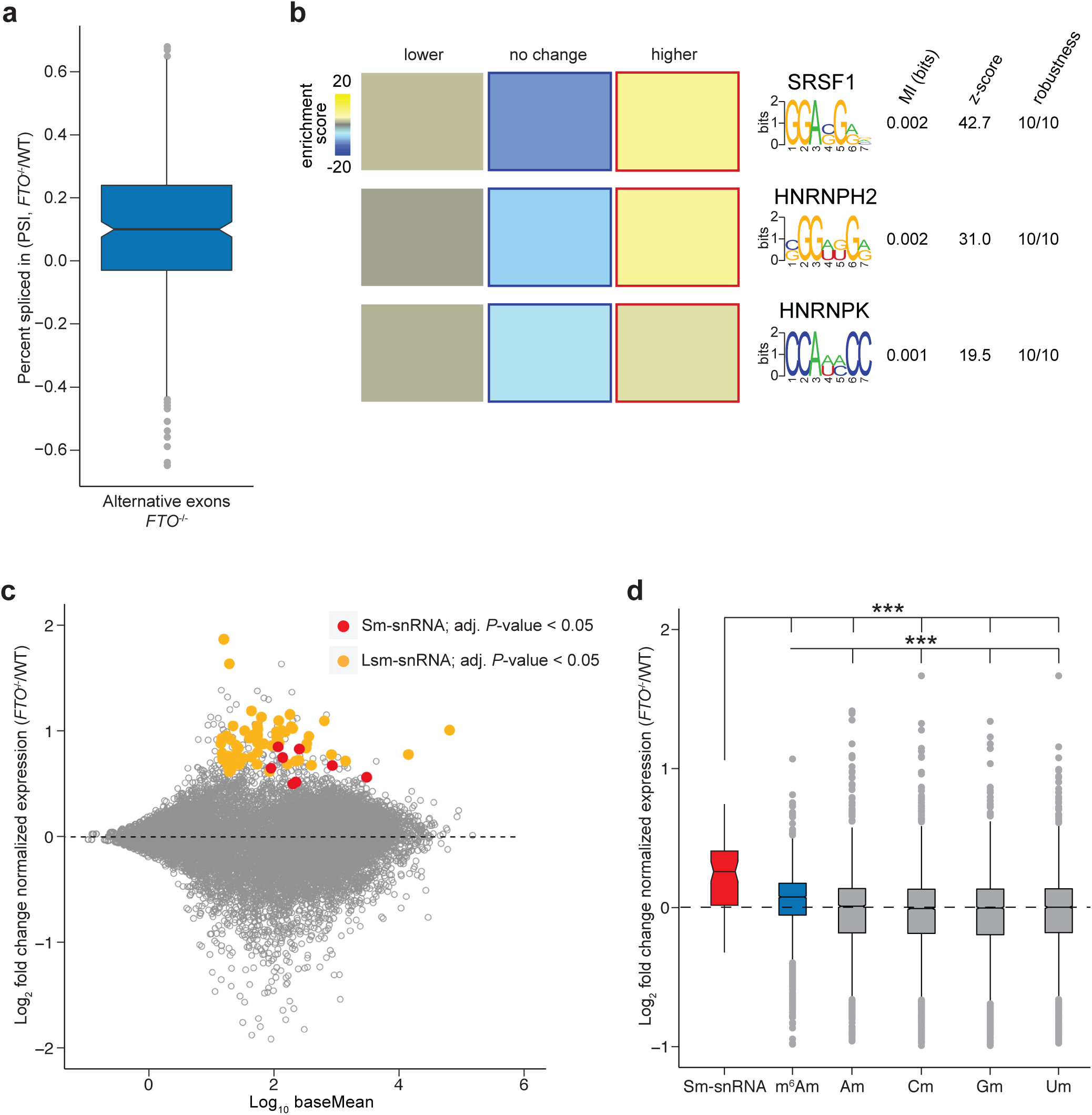
m2-snRNAs are highly abundant and promote SRSF1-dependent exon inclusion. **a,** FTO deficiency leads to exon inclusion. In line with previous reports^27,28^, transcriptome-wide alternative splicing analysis shows increased abundance of alternatively spliced exons in FTO deficient (*FTO*^-/-^) HEK293T cells. These data indicate that m_2_-snRNAs, which are the predominant snRNA isoform in FTO deficient cells, promote the inclusion of exons that are normally excluded in wild-type cells. (PSI = percent spliced in; Bayes factor cut-off > 10). **b,** m_2_-snRNA-regulated exons show enrichment of specific splicing regulator binding motifs. FIRE^53^ analysis revealed that the exons preferentially included in FTO-deficient HEK293T cells are enriched in binding sites for SRSF1, HNRNPH2 and HNRNPK. These data suggest that the presence of m_2_-snRNAs changes binding activity of specific splicing regulators. For each motif, we indicate the mutual information (MI) value, Z-score associated with the MI value, and the robustness score ranging from 0/10 to 10/10. The enrichment score indicates over-representation of a motif; significant over-representation is indicated by red frames, whereas blue frames indicate under-representation. **c**, FTO deficiency increases the abundance of Sm-class snRNAs. The MA plot of RNA-Seq analysis of total cellular RNA from previously published datasets^27^ shows the transcriptomewide log_2_ fold change of gene expression in *FTO* knockout HEK293T cells (*FTO*^-/-^) compared to wild-type HEK293T cells (WT). Significantly regulated Sm-class snRNAs (Sm-snRNAs) are indicated in red, significantly regulated Lsm-class snRNAs (Lsm-snRNAs) are indicated in orange. Notably, all significantly changed snRNAs show upregulation in *Fto*^*-/-*^cells. Since snRNAs largely exist in the m_2_-form in *FTO*^-/-^cells, these data suggest that m_2_-snRNAs are intrinsically more stable than m_1_-snRNAs that are predominately found in WT cells. Notably, Lsm-snRNAs (U6 and U6atac) were also upregulated in *Fto*^*-/-*^cells. Since Lsm-snRNAs are not directly regulated by FTO, these effects are likely to be indirect e.g. by changes in general snRNP turnover (Data represents the average of three independent biological replicates per genotype). **d,** m_2_-snRNAs show higher increase in expression levels than m^6^Am-initiated mRNAs in FTOdeficient cells. mRNAs were classified based on their annotated starting nucleotide (m^6^Am, 2’*O*-methyladenosine (Am), 2’-*O*-methylcytidine (Cm), 2’-*O*-methylguanosine (Gm) or 2’-*O*methyluridine (Um)). Shown are log_2_ fold changes of expression for snRNAs and mRNAs. As previously described^21^, FTO deficiency increases the abundance of m^6^Am-mRNAs compared to mRNAs starting with Am, Cm, Gm or Um. However, snRNAs were upregulated even more in FTO-deficient cells (FTO^-/-^) indicating that snRNAs are the major RNA class regulated by FTO under basal conditions. (Data represents the average from datasets of three independent biological replicates per genotype; one-way ANOVA with Tukey’s post hoc test \*\*\**P* ≤ 0.0001)

Recently, we found that methylation of Am to m^6^Am adjacent to mRNA caps confers substantially elevated stability to mRNA^21^. Thus, we asked if snRNAs show a similar increase in abundance when their m^6^Am levels are increased. To do this, we examined our own and preexisting^27^ RNA-Seq datasets of total cellular RNA from FTO-depleted cells. In both *FTO* knockout HEK293 cells and *Fto* knockout mouse liver, we observed an increase in snRNA levels relative to wild-type cells (**Figs. 4c, d** and **Extended Data Figs. 8c, d**). These data suggest that a function of m^6^Am is to increase snRNA levels in cells, which could contribute to the enhanced splicing activity associated with m_2_-snRNAs.

## Discussion

mRNA splicing involves snRNAs that are key components of spliceosomes, and regulatory proteins that influence the assembly and function of spliceosomes on pre-mRNAs. Although splicing regulation by trans-acting factors is well established, the modified nucleotides in snRNAs have not previously been thought to be the target of regulation. Here we show that snRNAs exist in two distinct methylation states which have distinct properties in cells. The methyl modification is located at the adenosine of the first encoded nucleotide in mRNA, is reversible, and influences snRNA abundance and splicing efficiency in cells. Together, our studies reveal that reversible epitranscriptomic information is encoded at the first nucleotide of snRNAs and determines their fate and function in cells.

The formation of m_2_-snRNAs and its subsequent demethylation to m_1_-snRNA is a previously unknown step in snRNA biogenesis (**Fig. 5**). snRNA biogenesis was extensively studied and involves Sm core assembly, 3’ processing, nucleotide modifications at internal sites and formation of the hypermethylated guanosine cap^6-8,11,14,17^. Our data reveal that spliceosomal snRNA biogenesis also involves a previously unrecognized *N*^6^-methylation that occurs on the evolutionarily conserved adenosine that is present at the transcription-start nucleotide of all Sm-class snRNAs. We find that FTO is a previously unknown snRNA biogenesis enzyme. FTO is involved in snRNA biogenesis by performing a regulated demethylation step of m_2_-snRNAs to m_1_-snRNAs. Thus, our studies reveal that snRNAs exhibit a more complex process of biogenesis than previously known, involving reversible methylation of the extended snRNA cap.

**Figure 5.**
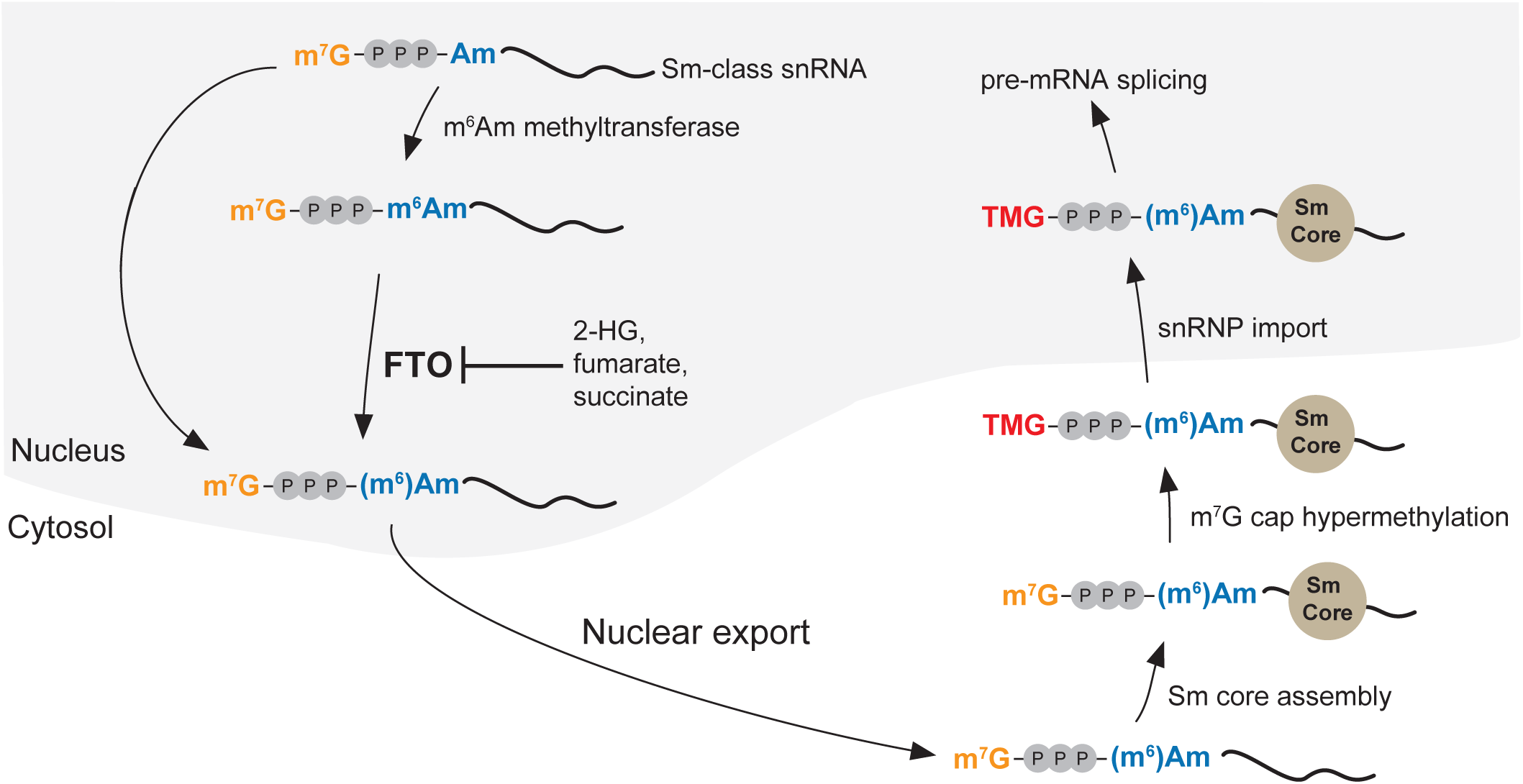
Proposed model of m_2_-snRNA biogenesis. Before their export to the cytosol, Sm-class snRNAs (m_1_-snRNAs) can be subjected to *N*^6^methylation of the m^7^G cap-adjacent 2’-*O*-methyladenosine (Am). This leads to the formation of m_2_-snRNAs containing an m^7^G cap-adjacent *N*^6^,2’-*O*-dimethyladenosine (m^6^Am). Under basal conditions, m_2_-snRNAs can be converted back to the m_1_ isoform by the nuclear RNA demethylase FTO. Similar to m_1_-snRNAs, m_2_-snRNAs are subjected to the normal biogenesis pathway, which involves Sm core assembly, hypermethylation of the m^7^G cap to *N*^2,2,7^-trimethylguanosine (TMG), reimport into the nucleus, and formation of functional small nuclear ribonucleoprotein complexes (snRNPs) that carry out pre-mRNA splicing. Notably, in conditions where specific metabolites, such as 2-hydroxyglutarate (2-HG), fumarate and succinate, are increased, FTO activity is inhibited^31,32^. This results in high levels of m_2_snRNAs, which leads to inclusion of alternative exons and ultimately determines transcriptome diversity.

Our finding that FTO is an snRNA biogenesis enzyme helps to explain earlier findings that FTO levels influence alternative splicing. These splicing changes were initially attributed to FTO-regulated m^6^A residues around splice-sites, since at the time FTO was thought to target m^6^A in mRNA based on its low, but measureable, demethylation activity towards m^6^A^27,28^. However, a role for m^6^A in alternative splicing has recently been challenged^38^. Our findings potentially provide a resolution to these discrepancies by showing that FTO regulates m^6^Am in spliceosomal snRNAs. Thus, our discovery of FTO-regulated m_2_-snRNAs provides the first direct link between FTO activity and the mRNA splicing machinery.

Notably, our transcriptome-wide analysis does not provide evidence for statistically significant alterations in m^6^A in any mRNA. Instead, our data reveals that FTO affects m^6^Am and nuclear RNAs, with snRNAs being robust targets of FTO. Our biochemical data shows that FTO can exhibit a pronounced and robust demethylation of m^6^Am in snRNA to near completion in many cell types. In contrast, depletion of FTO causes a clear and dramatic effect on the methylation state of snRNA, resulting in the majority of adenosine to be m^6^Am. This type of “methylation switch” between a methylated (m^6^Am) and demethylated (Am) nucleotide of RNA methylation is not seen at any m^6^Am or any m^6^A site in any mRNA to our knowledge, supporting the idea that the main physiologic targets of FTO are these small nuclear RNAs.

Studies of FTO localization show that FTO co-localizes with nuclear structures linked to premRNA splicing^19^, which is consistent with FTO’s action on these RNAs. Since snRNA biogenesis can cause diverse effects on the transcriptome, FTO depletion is expected to have diverse effects on mRNA. However, these effects likely reflect indirect effects of FTO on mRNA through its regulation of snRNA.

Notably, we used direct measurements of the specific methylated and demethylated nucleotides to determine how the stoichiometry of m^6^Am is controlled by FTO in snRNA. Measuring both the methylated and demethylated nucleotide at specific sites in the transcriptome is critical since it provides definitive evidence that FTO can regulate the stoichiometry of methylation. In contrast, immunoprecipitation of methylated RNA, followed by RNA quantification is problematic since the non-methylated nucleotide is not measured. Furthermore, changes in RNA immunoprecipitation can be caused by changes in the RNA levels in the input sample or by other methylated RNAs that compete for immunoprecipitation. Although our data support the idea that FTO – a nuclear-localized enzyme – primarily targets snRNAs, which are also primarily nuclear, it has been suggested that FTO can demethylate m^6^A in cytoplasmic mRNAs. However, as of yet, no m^6^A site has successfully been shown to be regulated by FTO using SCARLET, a quantitative method for determining m^6^A stoichiometries in mRNA^39^.

After snRNA transcription, the snRNA is methylated to the m_2_ form and then demethylated by FTO back to the m_1_ form (**Fig. 5**). The decision to demethylate the m_2_-snRNA appears to be determined early in snRNA biogenesis and may be important to establish splicing patterns in different cell types or metabolic states. We find that FTO-mediated demethylation of m_2_-snRNAs is inhibited when 2-hydroxyglutarate levels are increased due to expression of a cancer-associated mutant IDH. These data indicate that FTO is a metabolite sensor that connects metabolite abnormalities to snRNA methylation status. FTO was previously shown to be inhibited by metabolites that compete with *α*-ketoglutarate, including citrate, succinate, fumarate and 2-hydroxyglutarate^31,32^. Altered levels of these metabolites are seen in diverse cancers^40-43^, as well as during cellular differentiation^44,45^. Our studies raise the possibility that metabolite abnormalities may mediate their effects, in part, by inhibiting FTO and increasing the levels of m_2_-snRNAs that can in turn influence specific splicing events.

The presence of m_2_-snRNAs is not restricted to cells that lack FTO or express mutant IDH. We also detected m_2_-snRNAs in mouse embryonic stem cells, TF-1 cells and mouse liver tissue by miCLIP and mass spectrometry. It will be important to determine whether m_2_-snRNAs in these cells accumulate because of FTO inhibition or due to other mechanisms.

At present, the mechanism by which m^6^Am affects the function of snRNAs and snRNPs is not known. m^6^Am may recruit specific “readers,” may influence snRNA stability, or could have other effects. Notably, the levels of snRNAs are elevated in FTO-deficient cells. This includes the levels of U6 snRNA, an snRNA that lacks m^6^Am. The U6 snRNP is involved in complexes with other snRNPs that are dynamically assembled and disassembled during splicing. The global elevation in diverse snRNAs suggest that the overall pattern of snRNP turnover is affected by the presence of m_2_-snRNAs. However, a major next step will be to establish a mechanistic explanation for how the presence of m^6^Am influences snRNA/snRNP turnover and how this leads to altered splicing.

Notably, splicing efficiency is directly dependent on the levels of snRNAs and reduced snRNA expression is associated with severe splicing abnormalities^46^. Our data show that FTO deficiency leads to a clear and robust increase of snRNA expression. The increased snRNA expression may influence splicing patterns by specifically promoting the inclusion of exons that are otherwise inefficiently spliced due to rate-limiting levels of snRNPs.

It should be noted that the enhanced splicing activity of m_2_-snRNAs are unlikely to reflect a direct effect of the *N*^6^-methyl modification on the snRNA:pre-mRNA hybridization steps. These hybrids mostly occur at internal sites in the snRNA. For example, U1 snRNA hybridizes to the 5’ splice site using nucleotides 3-7 (ref. ^47^). Therefore, snRNA binding and recognition of target sites in pre-mRNA are unlikely to be promoted by the *N*^6^-methylation status of the first nucleotide.

Our data provide a new perspective on studies that have explored the physiological effects of FTO depletion. For example, FTO-deficient humans show developmental abnormalities and FTO-deficient mice show neuronal and metabolic defects^48^. Additionally, FTO-deficient leukemia cells show altered survival^49^, although other studies have not been able to replicate this effect^50^. Since snRNAs are the only individual transcripts that show statistically significant increases in our miCLIP analysis of *FTO* knockout cells, these phenotypes likely reflect the physiological consequences of aberrant activation of m_2_-snRNA-dependent splicing.

Although snRNAs were previously thought to be comprised of a single molecular species, the demonstration that snRNAs exist as distinct methyl isoforms indicates a potential regulatory role for methylation switches in splicing. Importantly, some snRNAs have non-splicing functions^30,51,52^ and the function of m_2_-snRNAs could therefore influence these other RNA processing pathways. More research will be needed to understand how the levels of m_2_-snRNPs are regulated and to identify the methyltransferase that introduces the *N*^6^-methyl modification in snRNAs.

## Acknowledgements

This work was supported by NIH grants R01DA037755 (S.R.J.), R01GM123977 (H.G.), R01NS102451 (L.P.), P01HD67244 and UO1HL121828 (S.S.G.), by the French Centre National de la Recherche Scientifique (T.G., J.-J.V., F.D.), DFG Priority Program grant RE2796/3-1 (A.R.) and by a DFG Research Fellowship (J.M.).

## Author Contributions

S.R.J., L.P. and J.M. designed the experiments. J.M. carried out the experiments. F.D., J.V. and T.G. synthesized modified oligonucleotides. A.R. produced hTGS1. S.S.G. and M.S. carried out mass spectrometry analysis. H.G. carried out the computational splicing analysis. S.R.J. and J.M. wrote the manuscript with input from all coauthors.

## Author Information

Correspondence and requests for materials should be addressed to S.R.J.

## Methods

### miCLIP-based mapping of 6mA at transcription start nucleotides and RNA-Seq analysis

For miCLIP experiments, total RNA was extracted from *Fto* knockout mouse livers (*Fto*^-/-^) and wild-type mouse livers using RNAzol RT (Molecular Research Center, Inc.). miCLIP was carried out as described previously^54^. In miCLIP, 6mA is mapped by immunoprecipitation and crosslinking with a 6mA-binding antibody (see below)^54^. This mapping method detects both m^6^A and m^6^Am, the two 6mA-containing nucleotides in RNA. RNA-Seq libraries of input and 6mA immunoprecipitated and crosslinked samples were prepared using a cloning strategy according to the miCLIP protocol^54,55^ and submitted to the Weill Cornell Medicine Epigenomics Core for sequencing on the Illumina HiSeq 2500 instrument, in paired-end mode, with 50 bases per read. Three independent biological replicates were sequenced for each condition. Raw input and miCLIP reads were trimmed of their 3’ adapter and demultiplexed as previously described^54^.

To analyze 6mA coverage at the transcription start nucleotide (TSN) the following steps were carried out. Input and miCLIP reads were aligned to mm10 with STAR to generate RPM-normalized bedgraphs and sorted bam files^56^ (version 2.5.2a;). Bedgraphs were visualized using the Integrative Genomics Viewer^57^ (IGV, version 2.4.4). Sorted bam files were converted to bed format and read coverage was determined in a 20-nucleotide window flanking annotated TSN (UCSC) using bedtools (version 2.25.0). miCLIP TSN coverage reads were then counted and normalized to input coverage reads using the Deseq2 pipeline adjusted for CLIP-Seq^58^ (version 1.18.1; commands: design= ∼ assay + condition + assay:condition; test=“LRT”, reduced= ∼ assay + condition). Analysis and visualization was carried out with custom in-house generated Rscripts using RStudio (Version 1.0.136). Only transcripts with normalized read counts > 10 were included in the analyses.

For all other RNA-Seq analyses, total RNA was diluted to a concentration of 50 ng/µl and submitted to the Weill Cornell Medicine Epigenomics Core for isolation of mRNA and library preparation using the Illumina TruSeq Stranded mRNA Library Prep Kit (RS-122-2101, Illumina). The libraries were sequenced on the Illumina HiSeq 2500 instrument, in paired-end mode, with 100 bases per read. At least two independent biological replicates were sequenced for each condition.

Previously published RNA-Seq datasets utilized in the current study^21,27^ were extracted from Gene Expression Omnibus (GEO, NCBI) and, if no processed data was available, the fastq-files were reanalyzed with the pipelines described above.

### Cell culture and animals

Flp-In T-REx HEK293 cells (R78007, ThermoFisher Scientific) were maintained in DMEM (ThermoFisher Scientific) with 10% FBS and antibiotics (100 units/ml penicillin and 100 µg/ml of streptomycin) under standard tissue culture conditions. Cells were split using TrypLE™ Express (Life Technologies) according to the manufacturer’s instructions.

Control, IDH1^R132H^ and IDH2^R140Q^ TF-1 erythroleukemia cells were generated and maintained as previously described^34^. Intracellular 2-hydroxyglutarate concentrations were determined as previously described^34^. All experiments involving TF-1 cells were carried out between passage 5-10. To inhibit production of neomorphic 2-hydroxyglutarate, IDH1^R132H^ cells or IDH2^R140Q^ cells were treated for 7 days with 1 µM AGI-5027 (SML1298, Sigma-Aldrich) or AGI-6780 (SML0895, Sigma-Aldrich), respectively.

Naive mouse embryonic stem cells (mESCs) were maintained in Knockout DMEM (ThermoFisher Scientific) with 15% FBS, antibiotics (100 units/ml penicillin and 100 µg/ml of streptomycin), 2mM L-glutamine, 50 µM *β*-mercaptoethanol (ThermoFisher Scientific), nonessential amino acids (ThermoFisher Scientific), recombinant mouse LIF (EMD Millipore), GSK3 inhibitor (04000402, Stemgent) and MEK1 inhibitor (04000602, Stemgent) under standard tissue culture conditions. To induce differentiation, naive mESCs were deprived of LIF, GSK3 inhibitor and MEK1 inhibitor for 7 days. Mycoplasma contamination in all cell lines used in this study was routinely tested by PCR.

*Fto* knockout mice were bred as previously described^24^. Only male mice at the age of 12-16 weeks were used for the experiments. All experiments involving mice were approved by the Institutional Animal Care and Use Committee at Weill Cornell Medicine.

### Antibodies

Antibodies used were as follows. For western blot analysis mouse *α*-FTO (ab92821, Abcam) and mouse *α*-ACTB (A2228, Sigma) were used. For 6mA-immunoprecipitation/miCLIP, rabbit *α*6mA (202-003, Synaptic Systems) was used. For SmB-IP, a previously described mouse *α*-SmB antibody (clone 18F6)^59^ was used. For 6mA immunoblotting, a rabbit *α*-6mA antibody (ab190886, Abcam) was used.

### Generation of *FTO* knockout cells

*FTO* knockout cells were generated by transfecting Double Nickase plasmids (sc-403708-NIC, Santa Cruz Biotechnology) containing two guide RNAs (Strand A: 5’CGGTCCCCTGGCCAGTGAAA-3’; Strand B: 5’-CCTGGTGTTCAGGTACTTGT-3’) into Flp-In T-REx HEK293 cells (Thermo Fisher Scientific) using LipoD293 (SignaGen Laboratories). 24 hours after transfection, GFP-positive cells were isolated by flow cytometry and reseeded. 48 hours after transfection, cells were subjected to puromycin selection (5 µg/ml) for three days. Cell were then reseeded at a density of 0.5 cells/well in 96-well plates for clonal selection. Loss of FTO protein expression was confirmed by Western blot. All experiments were carried out between passage 5-15.

### 6mA immunoblot

Denatured small RNA samples (5 µg) were separated by 10% polyacrylamide TBE-Urea denaturing gel electrophoresis at 200 V in 1x TBE. For blotting, separated RNA was transferred onto BrightStar-Plus positive charged nylon membranes (ThermoFisher Scientific) by semidry electroblotting in 0.5x TBE buffer at 300 mA for 30 min.

After UV crosslinking at 120,000 µJ/cm2 in a Stratalinker UV crosslinker (Stratagene), the membrane was blocked with 1% dry milk powder in PBS-T for 1 h at room temperature and then incubated with the *α*-6mA (1:500) overnight at 4°C. After washing with PBS-T, membranes were incubated with *α*-rabbit HRP-conjugated secondary antibody diluted 1:2500 with 1% dry milk powder in PBS-T for 1 h at room temperature. The membrane was visualized by Pierce ECL Western Blotting Substrate (ThermoFisher Scientific) and a Bio-Rad Gel Doc XR system. The quantification of the band intensity was analyzed using Bio-Rad Image Lab software.

### Determination of relative m^6^Am and Am levels in small RNAs by thin layer chromatography

The m^6^Am/Am ratio in small RNAs was determined as previously described^21^, with some modifications. The small RNA fraction (< 200 nucleotides) was isolated from total RNA using RNAzol RT (Molecular Research Center, Inc.).

For the analysis of individual U-RNAs, the small RNA fraction was separated on 6% TBE-Urea gels. Small RNAs were stained with SYBR Gold (ThermoFisher Scientific) and the respective bands (∼160 nucleotide for U1 snRNA, and ∼180 nucleotides for U2 snRNA) were excised and extracted using ZR small-RNA PAGE Recovery kit (Zymo Research). Notably, U1 snRNA comigrates with 5.8S rRNA. However, 5.8S rRNA is not a possible source of m^6^Am contamination, since it does not start with adenosine.

200 ng of the small RNA fraction or 40 ng of individual U-RNAs were decapped with 25 units of RppH (NEB) for 3h at 37 °C. The 5’ phosphates of the exposed cap-adjacent nucleotide were removed by the addition of 5 units rSAP (NEB) and further incubated for 1 hour at 37°C. Up to this point, all enzymatic reactions were performed in the presence of SUPERase In RNase Inhibitor (ThermoFisher Scientific). After phenol-chloroform extraction and ethanol precipitation, RNA samples were resuspended in 10 µl of DEPC-H_2_O and 5’ ends were labeled using 30 units T4 PNK and 0.8 mBq [γ-^32^P] ATP at 37°C for 30 min. PNK was heat inactivated at 65°C for 20 min and the reaction was passed through a P-30 spin column (Bio-Rad) to remove unincorporated isotope. The labeled RNA was then digested with 4 units of P1 nuclease (Sigma) for 3h at 37°C.

The nucleotide mixture was subsequently dried using an Eppendorf Vacufuge and reconstituted with 3 µL ddH_2_O. 1 µl of the released 5’ monophosphates from this digest were then analyzed by 2D-TLC on glass-backed PEI-cellulose plates (MerckMillipore) as described previously^21^. Developed TLC plates were imaged as described previously^21^.

### Protein expression and purification

Full-length, recombinant human FTO was expressed and purified as previously described^21^. Notably, to avoid any unspecific activity, it is advisable to express full-length FTO since the positively charged N-terminus of FTO may interact with the negatively charged m^7^G-ppp-cap of RNA.

Recombinant human TGS1 (hTGS1) was expressed and purified as previously described^60,61^. Briefly, *E. coli* Tuner cells transformed with pRSET-A-hTGS1_618-853_ were grown at 37 °C in 2YT medium until OD_600_ of 0.6 and induced with 0.2 mM IPTG. After overnight expression at 18°C, cells were lysed and the protein was purified via Ni-NTA (binding buffer: 50 mM Tris, pH 8, 1 M NaCl, 10 % glycerol; elution buffer: additional 500 mM imidazole) followed by gel filtration (Superdex 75, running buffer: 50 mM Tris, pH 7.5, 200 mM NaCl, 10 % glycerol) to obtain RNase-free protein that was concentrated to ∼1.5 mg/mL (MW: 27 kDa), flash frozen and stored at -20°C.

### Synthesis and characterization of synthetic oligonucleotides

Synthetic RNA oligonucleotides were chemically assembled as previously described^21^. To generate m^2,2,7^Gppp-capped oligonucleotides, m^7^Gppp-capped synthetic oligonucleotides (ONs) were treated with recombinant human TGS1. To do this, reaction mixtures containing 50 mM Tris-HCl (pH 8.0), 5 mM DTT, 50 mM NaCl, 1 mM AdoMet, and 150 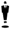 M pure m^7^G-ppp-ON (∼50 nmoles) and 2.5 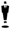 M hTgs1 were incubated at 37°C. Aliquots were withdrawn at different times and then analyzed by RP-HPLC to monitor the reaction (20 min linear gradient of 0-24% CH_3_CN in 50 mM TEAAc buffer pH 7). After 7 h incubation, the *N*2-Guanine-dimethylation was complete. Proteic material was then removed through a Sep Pak™ C_18_ cartridge. After dilution with 4 mL 100 mM TEAAc, pH 7 the crude mixture was loaded onto the cartridge and a wash was performed with 10 mL of 100 mM TEAAc then ON elution was made with 10 mL 50% CH_3_CN in 12.5 mM TEAAc. The solution was dried by lyophilization. Remaining AdoMet and the *S*-adenosylhomocysteine (AdoHcy) byproduct were removed as follows: the dry residue was dissolved with 1 mL H_2_O and the solution was loaded onto a Sephadex G-25 gel filtration cartridge (NAP™-10 cartridge). Elution was performed with 1 mL H_2_O. When necessary, m^2,2,7^Gppp-capped ONs were purified by IEX-HPLC and they were characterized by mass spectrometry MALDI-TOF as previously described^21^.

### Measurement of FTO activity

*In vitro* demethylation measurements were performed as described previously^21^. Briefly, demethylation activity assay was performed in 20 µl of reaction mixture containing 5 µM synthetic RNA oligonucleotide (either 5’-m^7^G-ppp-m^6^Am-CACUUGCUUUUGACACAACU-3’ or 5’-TMG-ppp-m^6^Am-CACUUGCUUUUGACACAACU-3’), 20 nM recombinant, full-length human FTO, 75 mM of (NH_4_)_2_Fe(SO_4_)_2_, 300 mM 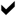-ketoglutarate, 2 mM sodium L-ascorbate, 150 mM KCl and 50 mM HEPES buffer, pH 7.0. The reaction was incubated at 37°C for 10 min and quenched by the addition of 1 mM of EDTA followed by inactivation of the enzyme for 5 min at 95°C.

### Sample preparation for HPLC analysis

After treatment with recombinant, full-length human FTO, oligonucleotides were decapped with 25 units of RppH (NEB) in ThermoPol buffer for 3 h at 37°C. RNA was subsequently digested to single nucleotides with 200 units S1 nuclease (Takara) for 2h at 37°C. 5’ phosphates were removed with 5 units rSAP (NEB) for 1h at 37°C. Before loading the samples onto the HPLC column, proteins were removed by size exclusion chromatography with a 10-kDa cut-off filter (VWR).

### HPLC analysis of demethylation activity

The HPLC analysis of nucleosides was performed on an Agilent 1100 system (Agilent Technologies). Separation was performed on a Poroshell 120 EC-C18 column (4µm, 150 × 4.6 mm, Agilent Technologies) equipped with an EC-C18 Guard cartridge (Agilent Technologies) at 22°C. The mobile phase consisted of buffer A (25 mM NaH_2_PO_4_) and buffer B (100% acetonitrile). Pump control and peak integration was achieved using the ChemStation software (Rev. A.10.02, build 1757, Agilent Technologies). Samples were analyzed at 2 ml/min flow rate with the following buffer A/B gradient: 7.5 min 95%/5%, 0.5 min 90%/10%, 2 min 10%/90%, 1 min 95%/5%. Retention times of the individual nucleosides were determined with synthetic standards (6.2 min for 2’-*O*-methyladenosine (Am), and 7.2 min for *N*^6^,2’-*O*-dimethyladenosine (m^6^Am).

### Sample preparation for mass spectrometry

In addition to their modified starting nucleotides (m^6^Am or Am), most snRNAs contain additional internal 2’-*O*-methylated adenosine (Am) residues^13^. Due to possible shearing of the RNA during the extraction process, these internal Am residues could be exposed and lead to unspecific background signal in thin layer chromatography. To eliminate this possible source of error, we sought to develop and independent method to measure the precise ratio of m^6^Am and Am caps in snRNA. We therefore developed a protocol to liberate intact cap-dinucleotides from the snRNA backbone and detect these cap-dinucleotides by LC-MS/MS.

To do this, cellular small RNA (0.1-1 µg) was digested with P1 nuclease (0.5-5 units) for 3 hours at 37°C. Notably, P1 nuclease does not cleave the triphosphate linker of the cap and thus specifically releases the cap-ppp-N dinucleotide, while digesting the RNA backbone down to single nucleotides. Proteins were removed by size exclusion chromatography with a 10-kDa cutoff filter (VWR).

The RNA was dried using an Eppendorf Vacufuge and reconstituted with 5 µL ddH_2_O. An aliquot was diluted 1:10-1:20 with 80% methanol in ddH_2_O. 2 µl of the resulting solution were subjected to LC-MS/MS analysis (see below).

### Identification of TMG-ppp-m^6^Am by mass spectrometry

An aliquot of the m^2,2,7^G-ppp-m^6^Am (TMG-ppp-m^6^Am) standard solution (generated by subjecting our synthetic 20-mer oligonucleotides to P1 nuclease digest) was subjected to LC/MSMS analysis by an Agilent 1200 LC-system coupled to an Agilent 6538 a quadrupole time-offlight mass spectrometer equipped with a dual electrospray ionization (ESI) source. A second isocratic pump delivered an internal reference mass solution (ions for negative mode 119.0360 and 966.0007; ions for positive mode 121.0509 and 922.0098) over the second nebulizer into the dual ESI source for continuous calibration during sample analysis. The sample injection volume was 4 µL. Chromatographic separation of the FTO-reaction products was performed using an aqueous normal phase (ANP) column (Cogent^™^ Diamond Hydride, 4 µm particle size, 150 mm x 2.1 mm; Microsolv Technology Corporation, NJ). To extend the column lifetime, a precolumn filter (0.5 µm, Microsolv) was placed in front of the ANP column. The mobile phase consisted of (A) 50% isopropanol with 0.025% acetic acid; and (B) 90% acetonitrile containing 5 mM ammonium acetate. To eliminate the interference of metal ions on the chromatographic peak integrity and ESI ionization, EDTA was added to the mobile phase in a final concentration of 6 µM. The final gradient applied was: 0-1.0 min 99% B, 1.0-7.0 min to 80% B, 7.0-18.0 min to 50% B, 18.0-19.0 min to 0% B and 19.1 to 33.0 min 99% B to regenerate the column. The flow rate was 0.4 mL/min. To reduce the amount of introduced salt and other interfering sample components, the first 0.5 min of the column effluent were directed to waste. Both positive and negative mass spectra were acquired in the 2 GHz extended dynamic range mode with 1.41 spectra/sec, sampled over a mass/charge range of 50-1000^62^. The operating ESI-source parameters for MS-analysis were: gas temperature 300 °C; drying gas flow 10 L/min; nebulizer pressure 35 psi; TOF capillary voltage 3500 V; fragmentor voltage 140V; skimmer voltage 65 V. Data was saved in centroid mode. Data was processed using Agilent MassHunter Qualitative Analysis Software (B.7. 00 Build 7.0.7024.0, Agilent Technologies).

### Analysis of TMG-ppp-m^6^Am and TMG-ppp-Am by mass spectrometry

Samples were injected into an LC/MS-system comprised of an Agilent 1260 HPLC and an Agilent 6460 triple quadrupole mass spectrometer (Agilent Technologies, Santa Clara, CA) equipped with a JetStream electrospray ionization source, using positive ion-monitoring in multiple reaction monitoring. The RNA-caps were resolved on an aqueous normal phase column (Cogent^™^ Diamond Hydride, 4 µm particle size, 150 mm x 2.1 mm; Microsolv Technology Corporation, NJ), at a column compartment temperature of 40°C. The samples were maintained at 4°C and the injection volume was 2 µL. The gradient-chromatography previously described by Chen *et al.*^62^ was optimized to achieve chromatographic separation of TMG-ppp-m^6^Am and TMG-ppp-Am. The aqueous mobile phase (A) was 50% isopropanol with 0.025% acetic acid, the organic mobile phase (B) was 90% acetonitrile containing 5 mM ammonium acetate. To eliminate the interference of metal ions on the chromatographic peak integrity and ESI ionization, EDTA was added to the mobile phase in a final concentration of 6 µM. The final gradient applied was: 0-1.0 min 99% B, 1.0-7.0 min to 80% B, 7.0-18.0 min to 50% B, 18.0-19.0 min to 0% B and 19.1 to 29.0 min 99% B to regenerate the column. The flow rate was 0.4 mL/min during data acquisition and 0.6 mL/min during re-equilibration from 21.0 to 28.5 min. Data was saved in centroid mode using Agilent Masshunter workstation acquisition software (B.06.00 Build 6.0.6025.4 SP4). Acquired raw data files were processed with Agilent MassHunter Qualitative Analysis Software (B.07.00 Build 7.0.7024.0, Agilent Technologies). The operating source parameters for MS-analysis were: gas temperature 300°C; gas flow 10 L/min; nebulizer pressure 35 psi; sheath gas temperature 400°C; sheath gas flow 11 L/min; capillary voltage 3500 V; nozzle voltage 0 V; fragmentor voltage 100V; cell accelerator voltage 7 V. Multiple reaction monitoring (MRM) data was acquired for the time segment 10-15 min in which the LC-flow was directed to the MS.

Product ion scans were performed at collision energies of 10, 20, 30, 40, 50 and 60 V, selecting [M+H]^+^ 843.2 as precursor ion for TMG-ppp-m^6^Am and [M+H]^+^ 829.1 for TMG-ppp-Am. Product ions were scanned in an *m/z* range of 50 – 850.

Optimized MRM transitions resulted at a collision energy of 50 eV and represented the deglycosylated base ions: for TMG-ppp-m^6^Am the transition 843.2→194.1* represented the formation of TMG and 843.2→150.1** the formation of m^6^Am, accordingly for TMG-ppp-Am 829.1→194.1* represented the formation of TMG and 829.1→136.1** the formation of Am. The transitions 843.2→438.1 and 829.1→424.1 were acquired as additional quality indicator. * indicates quantifier transitions, ** indicates the qualifier transitions.

### Immunoprecipitation of spliceosomal complexes

SmB immunoprecipitations were performed as described previously^36,37^. Briefly, extracts were prepared in ice cold RSB-100 buffer (100 mM NaCl, 10 mM Tris-HCl pH 7.4, 2.5 mM MgCl_2_) containing 0.1% NP40, protease, phosphatase and RNAse inhibitors, by passing five times through a 26-gauge needle followed by 3 × 5 seconds bursts of sonication on ice and centrifugation at 20,000 x *g* for 15 minutes at 4°C. Extract supernatant was quantified by BCA Assays (ThermoFisher Scientific).

For immunoprecipitation, a mouse monoclonal anti-SmB (18F6) antibody was bound to protein A/G-Sepharose (ThermoFisher Scientific) in RSB-100 buffer containing 0.1% NP40, protease, phosphatase and RNAse inhibitors for 2 hours at 4°C. Following five washes with the same buffer, antibody-bound beads were incubated with 200 µg of cell extract rotating for 2 hours at 4°C. Following five washes with the same buffer, samples were processed for RNA extraction.

Bound RNAs were extracted by treatment with proteinase K (200 µg) for 20 minutes at 37°C followed by TRIzol LS (ThermoFisher Scientific) extraction. Immunoprecipitated RNA was then analyzed by thin layer chromatography.

### Alternative splicing analysis

For alternative splicing analysis, paired reads were aligned to the human transcriptome (hg19) or mouse transcriptome (mm10) using STAR^56^ (v020201). Biological replicates were then merged and indexed using samtools (v1.5). The module exon_utils from the MISO package^63^ was first used to extract the constitutive exons from the transcriptome annotation (iGenomes, UCSC build hg19). The resulting annotations were then used in conjunction with the pe_utils module (MISO) to estimate fragment length distribution. The MISO package was then used to quantify percent spliced-in values (PSI) for each splicing event annotated for hg19 (pre-built MISO annotations). For this analysis, we used skipped exons (SE) as well as alternatively spliced first and last exons (A5SS and A3SS). We then used MISO to compare changes in splicing events, which reports the change in PSI values as well as the associated Bayes factor. For motif analysis, we extracted the included exons along with 250nt flanking sequences. We then used FIRE^53^, in non-discovery mode, to ask whether the presence or absence of known splicing regulator binding sites^64^ were significantly associated with changes in PSI values. In addition to enrichment profiles among the down- or up-regulated exons, FIRE also provides the mutual information (MI) values and their associated z-scores and robustness scores.

### Classification of mRNAs based on the first nucleotide

In experiments where we compared snRNAs to m^6^Am, Am, Cm, Gm, and Um-initiated mRNAs, we classified the mRNAs based on the nucleotide at the annotated transcription start site (TSS). Annotated TSS were extracted from UCSC table browser. A complete list of transcripts with their respective annotated transcription start site is found in the Supplementary Information Table S1.

### Code availability

All custom code used in this study can be obtained upon request from the lead author (S.R.J).

### Statistics and software

*P*-values were calculated with a two-tailed unpaired Student’s *t*-test or, for the comparison of more than two groups, with a one- or two-way ANOVA followed by Tukey’s post-test. *P*-values of 0.05 or less were considered significant.

---

**Extended Data Figure 1. FTO selectively demethylates small nuclear RNAs.**

**a**, Expression levels of FTO in *Fto*^*-/-*^liver and *FTO* knockout HEK293T cells. Upper panel shows a western blot analysis of FTO expression in wild-type (WT) and *Fto* knockout (*Fto*^-/-^) mouse liver samples that were used for 6mA miCLIP in **Fig. 1**. FTO protein was absent in *Fto*^*-/-*^livers. An antibody directed against *β*-ACTIN (ACTB) was used as a loading control. Lower panel shows a western blot analysis of FTO expression in wild-type (WT) and *FTO* knockout (*FTO*^-/-^) HEK293 cells that were used for the experiments in **Figs. 2, 3** and **4**. FTO protein was absent in *Fto*^*-/-*^HEK293 cells. An antibody directed against *β*-ACTIN (ACTB) was used as a loading control.

**b,** FTO deficiency leads to increased transcription-start nucleotide methylation of minor spliceosomal snRNAs. In **Fig. 1b**, we examined major spliceosomal snRNAs. Here we performed the same analysis on minor spliceosomal snRNAs. 6mA miCLIP reads were counted in a 20-nucleotide window surrounding the transcription-start nucleotide, normalized to input RNA-Seq in the same region and relative expression changes were calculated. In this analysis, no *P*-value cut-off was applied. Instead, the mean log_2_ fold change in transcription-start nucleotide methylation of specific snRNA gene classes (U11, U12, U4atac and U6atac) in *Fto* knockout mouse liver (*Fto*^-/-^) compared to wild-type liver (WT) is shown. Notably, Sm-class minor snRNA genes transcribed by RNA polymerase II increased methylation, whereas the U6atac snRNA gene, which is transcribed by RNA polymerase III and therefore does not acquire a m^7^G cap, shows no change in transcription-start nucleotide methylation upon FTO deficiency (Data represents the average from datasets of three independent biological replicates per genotype).

**Extended Data Figure 2. Read coverage tracks of minor spliceosomal small nuclear RNAs are markedly increased at transcription-start nucleotides in FTO-deficient cells.**

**a**, U11 snRNA shows increased transcription-start nucleotide methylation in FTO-deficient mouse liver. The grey tracks denote WT liver, whereas the blue track denotes *Fto*^*-/-*^liver. A representative read coverage track for U11 snRNA is shown. 6mA miCLIP reads are shown in the upper panel. Input reads are shown in the lower panel. The transcription-start nucleotide (TSN) is indicated in red (RPM = reads per million mapped reads; data represents the combined tracks from datasets of three independent biological replicates per genotype).

**b**, U12 snRNA shows increased transcription-start nucleotide methylation in FTO-deficient mouse liver. The grey tracks denote WT liver, whereas the blue track denotes *Fto*^*-/-*^liver. A representative read coverage track for U12 snRNA is shown. 6mA miCLIP reads are shown in the upper panel. Input reads are shown in the lower panel. The transcription-start nucleotide (TSN) is indicated in red (RPM = reads per million mapped reads; data represents the combined tracks from datasets of three independent biological replicates per genotype).

**c**, U4atac snRNA shows increased transcription-start nucleotide methylation in FTO-deficient mouse liver. The grey tracks denote WT liver, whereas the blue track denotes *Fto*^*-/-*^liver. A representative read coverage track for U4atac snRNA is shown. 6mA miCLIP reads are shown in the upper panel. Input reads are shown in the lower panel. The transcription-start nucleotide (TSN) is indicated in red (RPM = reads per million mapped reads; data represents the combined tracks from datasets of three independent biological replicates per genotype).

**d**, U6atac snRNA transcription-start nucleotide methylation is not affected in FTO-deficient mouse liver. The grey tracks denote WT liver, whereas the blue track denotes *Fto*^*-/-*^liver. A representative read coverage track for U6atac snRNA is shown. 6mA miCLIP reads are shown in the upper panel. Input reads are shown in the lower panel. The transcription-start nucleotide (TSN) is indicated in red (RPM = reads per million mapped reads; data represents the combined tracks from datasets of three independent biological replicates per genotype).

**Extended Data Figure 3. Read coverage tracks of U3 and U8 snoRNAs and U7 snRNA are increased at the transcription-start nucleotide in FTO-deficient cells.**

**a**, U3 snoRNA shows increased transcription-start nucleotide methylation in FTO-deficient mouse liver. The grey tracks denote WT liver, whereas the blue track denotes *Fto*^*-/-*^liver. A representative read coverage track for U3 snoRNA is shown. 6mA miCLIP reads are shown in the upper panel. Input reads are shown in the lower panel. The transcription-start nucleotide (TSN) is indicated in red (RPM = reads per million mapped reads; data represents the combined tracks from datasets of three independent biological replicates per genotype).

**b**, U7 snRNA shows increased transcription-start nucleotide methylation in FTO-deficient mouse liver. The grey tracks denote WT liver, whereas the blue track denotes *Fto*^*-/-*^liver. A representative read coverage track for U7 snRNA is shown. 6mA miCLIP reads are shown in the upper panel. Input reads are shown in the lower panel. The transcription-start nucleotide (TSN) is indicated in red (RPM = reads per million mapped reads; data represents the combined tracks from datasets of three independent biological replicates per genotype).

**c**, U8 snoRNA shows increased transcription-start nucleotide methylation in FTO-deficient mouse liver. The grey tracks denote WT liver, whereas the blue track denotes *Fto*^-/-^ liver. A representative read coverage track for U8 snoRNA is shown. 6mA miCLIP reads are shown in the upper panel. Input reads are shown in the lower panel. The transcription-start nucleotide (TSN) is indicated in red (RPM = reads per million mapped reads; data represents the combined tracks from datasets of three independent biological replicates per genotype).

**Extended Data Figure 4. *N*^6^,2’-*O*-dimethyladenosine (m^6^Am) is present in U2 snRNAs.**

**a**, FTO deficiency leads to a marked increased in *N*^6^,2’-*O*-dimethyladenosine (m^6^Am) in U2 snRNA. The relative abundance of modified adenosines at the transcription-start nucleotide in U2 snRNA caps derived from wild-type (WT) and FTO-deficient (*FTO*^-/-^) HEK293 cells was determined by thin layer chromatography. The left panel shows a TBE-Urea gel image stained with SYBR Gold, where the red dashed line indicates the U2-enriched fraction that was used for the analysis. The migration position of m^6^Am and 2’-*O*-methyladenosine (Am) is indicated by the dashed black circles. Additional spots are seen on the TLC chromatogram due to RNA fragmentation that occurs during RNA purification, resulting in 5’ radiolabeling at internal sites^23^.

The right panel shows the quantification of the m^6^Am/Am ratio in the U2-enriched fraction (*n* = 3 independent biological replicates; mean ± s.d; unpaired student’s *t*-test \*\**P* ≤ 0.001).

**Extended Data Figure 5. Development of an LC-MS/MS-based detection method for snRNA caps.**

**a**, Schematic of sample preparation for LC-MS/MS analysis. RNA is digested with P1, which cleaves after each nucleotide and releases 5’-monophosphorylated nucleotides (p-N). However, since the triphosphate linker between the RNA cap and the first nucleotide is resistant to P1 nuclease cleavage, the intact extended cap is released as a cap-dinucleotide (p = phosphate; N = nucleotide; P1 = P1 nuclease; grey arrows indicate P1 cleavage sites).

**b**, Positive ion mode fragmentation at 50 V collision energy of a TMG-ppp-m^6^Am and a TMGppp-Am standard solution. Left panel: total ion current (TIC) of the product ion scan (m/z 50-850) of a (A) TMG-ppp-Am (precursor ion [M+H]^+^ 843.1 and a (B) TMG-ppp-m^6^Am (precursor ion [M+H]^+^ 829.1) standard solution. Right panel: MS/MS product ion spectra of the integrated areas of the peaks for (A) and (B). Chemical structures indicate the most prominent product ions for (A) 135.8 ([adenine+H]^+^), 194.0 ([trimethylguanine+H]^+^) and 423.6 ([2’-*O*-methyladenosine-pp+H]^+^); for (B) 150.1 ([*N*^6^-methyladenine+H]^+^), 193.9 ([trimethylguanine+H]^+^) and 438.1 ([*N*^6^,2’-*O*-dimethyladenosine-pp+H]^+^).

**c**, Standard curves for the ratio of the MRM transitions 843.2 → 194.1 TMG-ppp-m^6^Am to 829.1 → 194.1 for TMG-ppp-Am (MRM) recorded in positive ion mode. Standard dilutions (0.006-4 ng/µL) of TMG-ppp-m^6^Am (stock solution contained 400 ng/µl overall nucleotides in water, measured by nano-drop analysis) in a small RNA extract of wildtype HEK-cells (100 ng/µl overall nucleotide concentration in 80% methanol) in 80% methanol were subjected to LCMS/MS analysis (n = 3, technical replicates). (left panel) A linear relationship of the MRM ratio of TMG-ppp-m^6^Am to TMG-ppp-Am over a nucleotide concentration range of 0.0125 to 4 ng/µl was determined. (right panel) After linear regression over a concentration range of 0.05 to 4 ng/µl with an R^2^ of 0.9987, a second dilution series of 0.006 to 0.10 ng/µl was analyzed to determine an LOQ of 0.0125 ng/µl.

**Extended Data Figure 6. m_2_-snRNAs are differentially regulated in different cells and conditions.**

**a**, FTO deficiency increases the relative abundance of m_2_-snRNA caps in HEK293 cells. Related to **Fig. 2c**: Small RNA from WT and *Fto*^*-/-*^was digested with nuclease P1 to specifically liberate the extended cap structure dinucleotide and analyzed by LC-MS/MS. Shown is the abundance of m_2_-snRNA caps (m^6^Am) and m_1_-snRNA caps (Am) represented by the integrated peak area of the corresponding MRM transitions (recorded in positive ion mode) (n=3 independent biological replicates, mean ± s.d.; unpaired Student’s t-test, *** p < 0.001).

**b**, Differentiation increases relative abundance of m_2_-snRNA (cap-ppp-m^6^Am) caps in mouse embryonic stem cells (mESCs). Small RNA from naïve and differentiated (diff.) mESCs was digested with nuclease P1 to specifically liberate the extended cap structure dinucleotide and analyzed by LC-MS/MS. Shown is the ratio of m_2_-snRNA caps (m^6^Am) to m_1_-snRNA caps (Am) represented by the integrated peak area ratio of the corresponding MRM transitions (recorded in positive ion mode). These data suggest that mESC differentiation induces a specific program that changes the abundance of m_2_-snRNAs (n=3 independent biological replicates, mean ± s.d.; unpaired Student’s t-test, ** p < 0.01).

**c**, Differentiation increases relative abundance of m_2_-snRNA (cap-ppp-m^6^Am) caps in mouse embryonic stem cells (mESCs). Related to **Extended Data Fig. 6b**: Small RNA from naïve and differentiated (diff.) mESCs was digested with nuclease P1 to specifically liberate the extended cap structure dinucleotide and analyzed by LC-MS/MS. Shown is the abundance of m_2_-snRNA caps (m^6^Am) and m_1_-snRNA caps (Am) represented by the integrated peak area of the corresponding MRM transitions (recorded in positive ion mode). These data suggest that mESC differentiation induces a specific program that changes the abundance of m_2_-snRNAs (n=3 independent biological replicates, mean ± s.d.; unpaired Student’s t-test, ** p < 0.01).

**d**, m_2_-snRNA are not detected in wild-type mouse liver and brain. Related to **Figs. 2d** and **3b**: Small RNA from wild-type (WT) and *Fto* knockout (*Fto*^-/-^) mouse liver and brain tissue was digested with nuclease P1 to specifically liberate the extended cap structure dinucleotide and analyzed by LC-MS/MS. Shown is the abundance of m_2_-snRNA caps (m^6^Am) and m_1_-snRNA caps (Am) represented by the integrated peak area of the MRM transition (recorded in positive ion mode). (n.d.=not detected; n=3 independent biological replicates, mean ± s.d.; unpaired Student’s t-test, ** p < 0.01).

**Extended Data Figure 7. m2-snRNAs are reversibly regulated in oncometabolite-dependent cancer models.**

**a**, Intracellular 2-hydroxyglutarate (2-HG) concentrations in mutant IDH1^R132H^ and IDH2^R140Q^ TF-1 cells (*n* = 3 independent biological replicates; mean ± s.d; one-way ANOVA with Tukey’s post hoc test \*\**P* ≤ 0.001).

**b**, Increased abundance of m_2_-snRNA caps in oncometabolite-dependent cancer. Related to **Fig. 2e**: Small RNA from Ctrl and mutant IDH-expressing cells was digested with nuclease P1 to specifically liberate the extended cap structure dinucleotide and analyzed by LC-MS/MS. Shown is the abundance of m_2_-snRNA caps (m^6^Am) and m_1_-snRNA caps (Am) represented by the integrated peak area of the cap-specific MRM transitions (recorded in positive ion mode). IDH1^R132H^and IDH2^R140Q^-expressing TF-1 cells have high levels of 2-hydroxyglutarate (2-HG). 2-HG is a natural inhibitor of FTO activity^32^ and leads to increased abundance of m_2_-snRNAs. Specific inhibition of the mutant IDH1 (AGI-5027) and IDH2 (AGI-6780) isoforms shows that these effects are readily reversible (n=3 independent biological replicates, mean ± s.d.; one-way ANOVA with Tukey’s post hoc test \**P* ≤ 0.05, \*\**P* ≤ 0.01).

**c**, Increased abundance of m^6^Am caps in U1 snRNA of oncometabolite-dependent cancer cells. Gel-extracted U1 snRNA from Ctrl and mutant IDH2^R140Q^ TF-1 cells was digested with nuclease P1 to specifically liberate the extended cap structure dinucleotide and TLC. Shown is the ratio of cap-adjacent m^6^Am and cap-adjacent Am. IDH2^R140Q^-expressing TF-1 cells have high levels of 2-hydroxyglutarate (2-HG). 2-HG is a natural inhibitor of FTO activity and leads to increased abundance of m^6^Am caps in snRNA. Specific inhibition of the mutant IDH2^R140Q^ isoform with AGI-6780 shows that these effects are readily reversible (n=3 independent biological replicates, mean ± s.d.; unpaired Student’s t-test, *** p < 0.001).

**Extended Data Figure 8. m2-snRNAs are highly abundant and promote SRSF1-dependent exon inclusion.**

**a,** Splicing regulator binding motif-enrichment in m_2_-snRNA-regulated exons. Related to **Fig. 4b**: FIRE^53^ analysis of exons preferentially included in FTO-deficient HEK293 cells. Binding motifs for U2AF2, HNRNPA1, HNRNPA2B1, SRSF2 and SRSF10 are not enriched in the included exons. These data suggest that changes of splicing regulator binding activity in the presence of m_2_-snRNAs are restricted to SRSF1, HNRNPH2 and HNRNPK, which promotes specific splicing patterns (see **Fig. 4b**). For each motif, we indicate the mutual information (MI) value, Z-score associated with the MI value, and the robustness score ranging from 0/10 to 10/10. The enrichment score indicates over-representation of a motif; significant over-representation is indicated by red frames, whereas blue frames indicate under-representation.

**b,** m_2_-snRNA-regulated exons show enrichment of specific splicing regulator binding motifs. To verify the experiments presented in **Fig. 4b** and **Extended Data Fig. 8a**), we performed FIRE^53^ analysis on exons included in an independent FTO knockdown from HEK293T cells. We observed a similar specific enrichment of SRSF1, HNRNPH2 and HNRNPK motifs as in FTO knockout cells. For each motif, we indicate the mutual information (MI) value, Z-score associated with the MI value, and the robustness score ranging from 0/10 to 10/10. The enrichment score indicates over-representation of a motif; significant over-representation is indicated by red frames, whereas blue frames indicate under-representation.

**c**, FTO deficiency increases the abundance of snRNAs. The MA plot shows the transcriptome-wide log_2_ fold change of gene expression in *Fto* knockout mouse liver (*Fto*^-/-^) compared to wildtype (WT) liver. Significantly regulated snRNAs are indicated in red. Notably, the significantly changed snRNAs were exclusively upregulated in *Fto*^-/-^ liver. Since snRNAs largely exist in the m_2_-form in *Fto*^*-/-*^mice, these data suggest that m_2_-snRNAs are intrinsically more stable than m_1_snRNAs that are predominately found in WT mice (Data represents the average from datasets of three independent biological replicates per genotype).

**d,** snRNA expression is broadly increased upon FTO deficiency in mice. Shown are log_2_ fold changes of snRNA expression in FTO knockout (*Fto*^-/-^) mouse liver. These data suggest that the outcome of FTO deficiency i.e. elevated snRNA expression is conserved between humans and mice. (Data represents the average from datasets of three independent biological replicates per genotype)

